# Decoding the development of the blood and immune systems during human fetal liver haematopoiesis

**DOI:** 10.1101/654210

**Authors:** Dorin-Mirel Popescu, Rachel A. Botting, Emily Stephenson, Kile Green, Laura Jardine, Emily F. Calderbank, Mirjana Efremova, Meghan Acres, Daniel Maunder, Peter Vegh, Issac Goh, Yorick Gitton, Jongeun Park, Krzysztof Polanski, Roser Vento-Tormo, Zhichao Miao, Rachel Rowell, David McDonald, James Fletcher, David Dixon, Elizabeth Poyner, Gary Reynolds, Michael Mather, Corina Moldovan, Lira Mamanova, Frankie Greig, Matthew Young, Kerstin Meyer, Steven Lisgo, Jaume Bacardit, Andrew Fuller, Ben Millar, Barbara Innes, Susan Lindsay, Michael J. T. Stubbington, Monika S. Kowalczyk, Bo Li, Orr Ashenbrg, Marcin Tabaka, Danielle Dionne, Timothy L. Tickle, Michal Slyper, Orit Rozenblatt-Rosen, Andrew Filby, Alexandra-Chloe Villani, Anindita Roy, Aviv Regev, Alain Chedotal, Irene Roberts, Berthold Göttgens, Elisa Laurenti, Sam Behjati, Sarah A. Teichmann, Muzlifah Haniffa

**Affiliations:** Institute of Cellular Medicine, Newcastle University, Newcastle upon Tyne, NE2 4HH, UK; Department of Haematology and Wellcome and MRC Cambridge Stem Cell Institute, University of Cambridge, Cambridge, CB2 2XY, UK; Wellcome Sanger Institute, Wellcome Genome Campus, Hinxton, Cambridge CB10 1SA, UK; European Molecular Biology Laboratory, European Bioinformatics Institute (EMBL-EBI), Wellcome Genome Campus, Cambridge, CB10 1SD UK; Sorbonne Université, INSERM, CNRS, Institut de la Vision, 17 Rue Moreau, F-75012 Paris, France; Department of Pathology, Newcastle Hospitals NHS Foundation Trust, Newcastle upon Tyne NE2 4LP, UK; Institute of Genetic Medicine, Newcastle University, Newcastle upon Tyne, NE1 3BZ, UK; School of Computing, Newcastle University, NE4 5TG, UK; Broad Institute of Harvard and MIT, Cambridge, MA 02142, USA; Center for Immunology and Inflammatory Diseases, Massachusetts General Hospital, Boston, MA 02129, USA; Klarman Cell Observatory, Broad Institute of Harvard and MIT, Cambridge, MA, USA; Celsius Therapeutics, Cambridge, USA; Data Sciences Platform, Broad Institute of Harvard and MIT, Cambridge, MA, USA; Department of Paediatrics, University of Oxford, Oxford OX3 9DS, UK; Howard Hughes Medical Institute, Koch Institute of Integrative Cancer Research, Department of Biology, Massachusetts Institute of Technology, Cambridge, MA, USA; MRC Molecular Haematology Unit and Department of Paediatrics, Weatherall Institute of Molecular Medicine, University of Oxford, and BRC Blood Theme, NIHR Oxford Biomedical Centre, Oxford OX3 9DS, UK; Department of Paediatrics, University of Cambridge, Cambridge CB2 0SP, UK; Theory of Condensed Matter Group, Cavendish Laboratory/Department of Physics, University of Cambridge, Cambridge CB3 0HE, UK; Department of Dermatology and NIHR Newcastle Biomedical Research Centre, Newcastle Hospitals NHS Foundation Trust, Newcastle upon Tyne NE2 4LP, UK

**Keywords:** Human development, haematopoiesis, immunology, single cell RNA-sequencing, liver, skin, kidney

## Abstract

Definitive haematopoiesis in the fetal liver supports self-renewal and differentiation of haematopoietic stem cells/multipotent progenitors (HSC/MPPs), yet remains poorly defined in humans. Using single cell transcriptome profiling of ~133,000 fetal liver and ~65,000 fetal skin and kidney cells, we identify the repertoire of blood and immune cells in first and early second trimesters of development. From this data, we infer differentiation trajectories from HSC/MPPs, and evaluate the impact of tissue microenvironment on blood and immune cell development. We predict coupling of mast cell differentiation with erythro-megakaryopoiesis and identify physiological erythropoiesis in fetal skin. We demonstrate a shift in fetal liver haematopoietic composition during gestation away from being erythroid-predominant, accompanied by a parallel change in HSC/MPP differentiation potential, which we functionally validate. Our integrated map of fetal liver haematopoiesis provides a blueprint for the study of paediatric blood and immune disorders, and a valuable reference for understanding and harnessing the therapeutic potential of HSC/MPPs.

## Introduction

The human blood and immune systems develop during early embryogenesis, and are critical for the survival and health of the organism. To date, our understanding of human haematopoietic development derives primarily from murine and *in vitro* model systems due to difficulties in human fetal tissue access and experimental manipulation. While haematopoietic development is conserved across vertebrates^1^, important differences between mouse and human have been noted^2,3^. For example, the dominance of fetal liver haematopoiesis until birth in mice^4^, and the earlier establishment of fetal bone marrow (BM) dominance in humans^5^. Therefore, interrogation of human tissue is vital to understand the molecular and cellular landscape of the developing blood and immune system.

The implications of fetal haematopoiesis reach beyond life *in utero* to physiological and pathological states in children and adults, and are relevant to therapeutic use of haematopoietic stem cells (HSC/MPPs) and to regenerative medicine. Further characterisation of fetal HSC/MPPs will inform inducible pluripotent stem cell technology and interventions to ‘supercharge’ cord blood or adult HSC/MPPs for clinical therapy. A molecular map of the developing blood and immune system will highlight potential mechanisms of monogenic primary immunodeficiencies, and childhood leukaemias^6^ and anaemias which originate in fetal life^7^.

The earliest blood and immune cells originate outside the embryo, arising from the yolk-sac between 2-3 post-conception weeks (PCW). At 3-4 PCW, intra-embryonic progenitors from the aorta-gonad-mesonephros (AGM) develop^8–10^. Yolk-sac and AGM progenitors colonise fetal tissues such as the liver, which remains the major organ of haematopoiesis until the mid-second trimester. Fetal BM is colonised around 11 PCW and becomes the dominant site of haematopoiesis after 20 PCW in human^8–10^. Yolk sac-, AGM-, fetal liver- and BM-derived immune cells seed peripheral tissues including non-lymphoid tissues (NLTs), where they undergo specific maturation programs which are both intrinsically determined and extrinsically nurtured by the tissue microenvironment^11,12^. Systematic, comprehensive analysis of multiple blood and immune lineages during human development has not previously been attempted.

In this study, we used single cell transcriptomics to map the molecular states of human fetal liver cells between 6-18 PCW, when the liver represents the predominant site of human fetal haematopoiesis. We integrate imaging mass cytometry, flow cytometry and cellular morphology to validate the transcriptome-based cellular profiles. We construct the functional organisation of the developing immune network through comparative analysis of immune cells in fetal liver with those in skin and kidney as representative non-lymphoid tissues (NLT).

## Results

### Single cell transcriptome map of fetal liver

To investigate blood and immune cell development in the fetal liver, we generated single cell suspensions from embryonic and fetal livers between 6 and 18 PCW. We FACS-isolated cells for droplet-based (10x Genomics) and indexed plate-based (Smart-seq2) single cell RNA sequencing (scRNA-seq) (Figure 1a-b, Extended Data Table 1, see Methods). Immunostaining with CD45 (10x) and CD45 and HLA-DR (Smart-seq2) allowed enrichment of cells within the respective gates (Supplementary Figure 1a). Where possible, we analysed skin and kidney cells from the same fetuses at the key initiation and amplification phase of fetal liver haematopoiesis between 6-12 PCW. This permitted parallel evaluation of blood and immune cell topography in non-lymphoid tissues (NLT) (Figure 1a-b).

**Figure 1:**
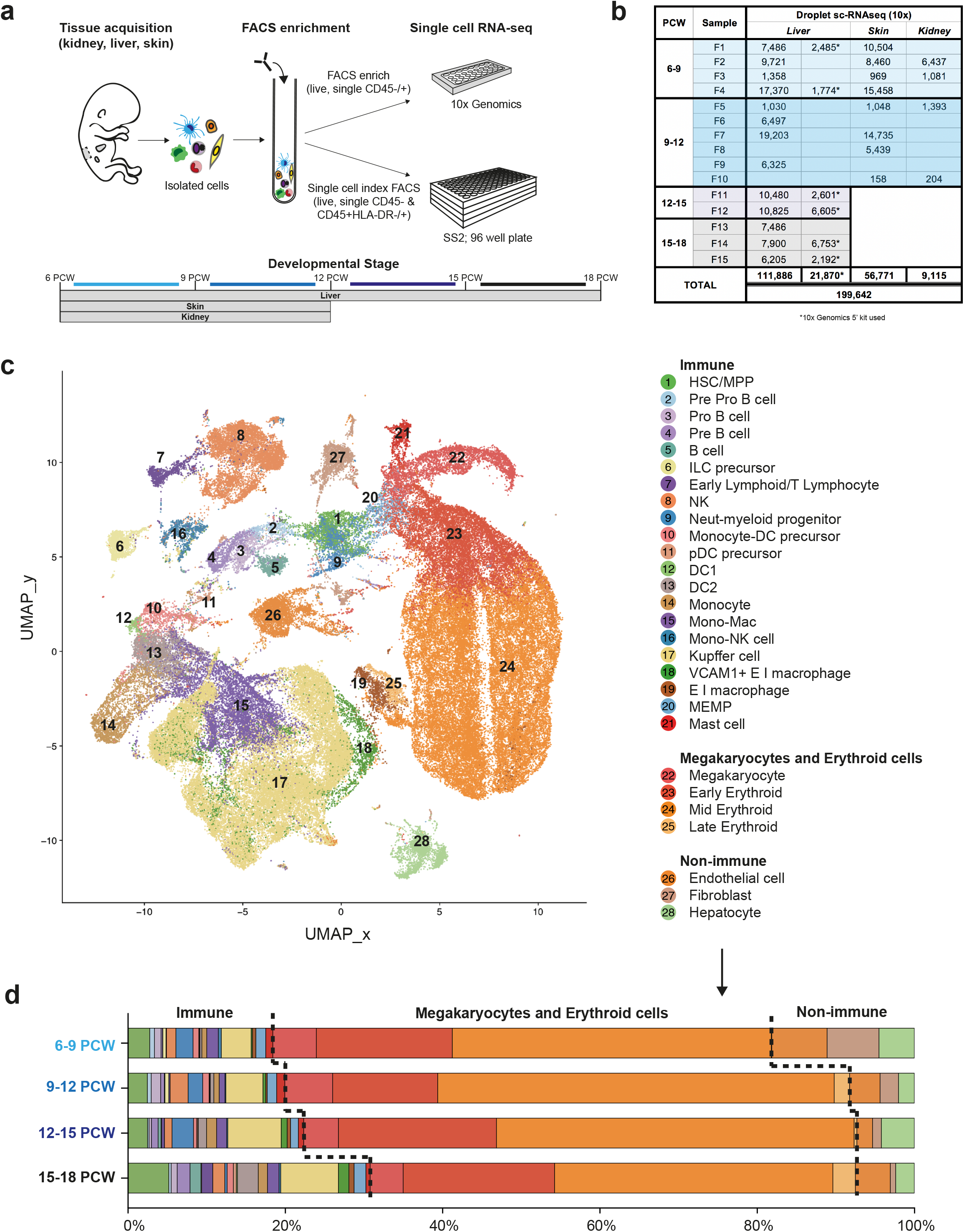
Single cell transcriptome map of fetal liver. **a**, Schematic of tissue processing and cell isolation for scRNA-seq. **b**, Number of fetal liver, skin and kidney cells (n=15) sequenced using 10x Genomics platform that passed QC. **c**, UMAP visualisation of 111,886 fetal liver cells with annotations. **d**, Mean % of each cell type from **c** over four developmental stages, corrected based on the proportion of cells in the initial sort gate (CD45^+^/CD45^−^) the subset was found in. SS2 = Smart-seq2; scRNA-seq = single cell RNA sequencing; HSC/MPP = haematopoietic stem cell/multipotent progenitor; ILC = innate lymphoid cell; NK = natural killer; Neut-myeloid = neutrophil myeloid; Monocyte-DC = monocyte-dendritic cell; pDC = plasmacytoid dendritic cell; DC = dendritic cell; Mono-mac = monocyte-macrophage; Mono-NK = monocyte-natural killer EI = erythroblastic island; MEMP = megakaryocyte-erythroid-mast cell progenitor; PCW = post conception weeks

In total, we performed scRNA-seq of 145,040 cells from liver of which 133,756 passed QC (111,886 cells using 3’kit and 21,870 cells using 5’ V(D)J kit; n = 13), 56,800 cells from skin with 56,771 passing QC (n = 8), and 9,171 cells from kidney with 9,115 passing QC (n = 4) (Figure 1b and Extended Data Table 2). An additional 3,571 fetal liver, skin and kidney cells from 9-12 PCW were profiled using the Smart-seq2 protocol (Extended Data Table 1). We detected on average ~3,000 genes per cell with the 10x Genomics platform and ~6,000 genes with the Smart-seq2 protocol (see Methods). We excluded cells with <200 genes, >20% mitochondrial gene expression and those identified as doublets (see Methods).

Our fetal liver dataset comprises 13 karyotypically normal fetuses: seven female and six male (Extended Data Figure 1a). The presence of maternal cells was excluded using genotyping from mRNA reads (see Methods). We performed graph-based Louvain clustering on 111,886 single cell transcriptomes generated by 10x 3’kit and derived marker genes to annotate the cell clusters (see Methods, Figure 1c). Consistency and agreement of cell clusters was additionally evaluated with a non-graph-based clustering method (Gaussian mixture) and a second graph-based (Agglomerative with Ward linkage) clustering method (Extended Data Figure 1b). Batch alignment was performed using Canonical Correlation Analysis^13^ and was visualised by UMAP. 28 major cell types were identified in the fetal liver (Figure 1c). We annotated cell types by cross-referencing cluster-defining transcripts with published literature reports of these genes and their cell-type specific expression (see Methods). We applied a descriptive nomenclature for these cell types based on their gene expression profile.

To evaluate fetal liver cell populations across developmental stages, we defined four gestational phases spanning the first and second trimesters: 6-9 PCW (40,194 cells; n = 4), 9-12 PCW (33,055 cells; n = 4), 12-15 PCW (30,775 cells, n=2), and 15-18 PCW (29,514 cells, n=3) (Figure 1b), and calculated the relative frequency of cell states during each phase (Figure 1d). All 28 cell states are present throughout these stages (Figure 1d and Extended Data Figure 1c) with 26/28 cell states represented in the sparser Smart-seq2 dataset (Extended Data Figure 1d). We did not detect neutrophils, basophils or eosinophils in the fetal liver, which is consistent with reports of granulocytes developing later during fetal BM haematopoiesis^14^. Our findings agree with previous reports on erythroid lineage bias in early fetal liver, and an increasing prominence of lymphoid and myeloid lineages with advancing gestation (Figure 1d and Extended Data Figure 1 e-f)^5^.

We provide a comprehensive expression profile of genes known to cause primary immunodeficiencies^15^ in the 28 inferred fetal liver cell types. The expression patterns of these genes will provide insight on the stage of onset of the penetrance of these mutations, and aid future molecular phenotyping studies of these disorders (Supplementary Figure 2).

### Validation of markers and cell states

We derived 150 cluster-defining marker genes (Extended Data Table 3) using the Seurat package (see Methods), which we manually trimmed to 48 genes (Figure 2a), where the limited set is still capable of discriminating all 28 cell states. To assess the relative performance of the 150-gene versus 48-gene list to classify cells, we used the ensemble learning random forest method, which gave 92% and 88% average precision for their respective ability to identify cells (Figure 2b). We selected genes encoding surface proteins (denoted by * in Figure 2a) to devise a FACS panel for cell isolation corresponding to the scRNA-seq defined clusters (Supplementary Figure 1b). We validated 18 fetal liver cell types by bulk transcriptome profiling from the respective FACS gates (Supplementary Figure 1b and 3a). Cytospin preparations of populations from these gates demonstrated morphology consistent with their designated cell identity, reinforcing the validity of our annotation and utility of single-cell transcriptomics to resolve cellular heterogeneity (Figure 2c).

**Figure 2:**
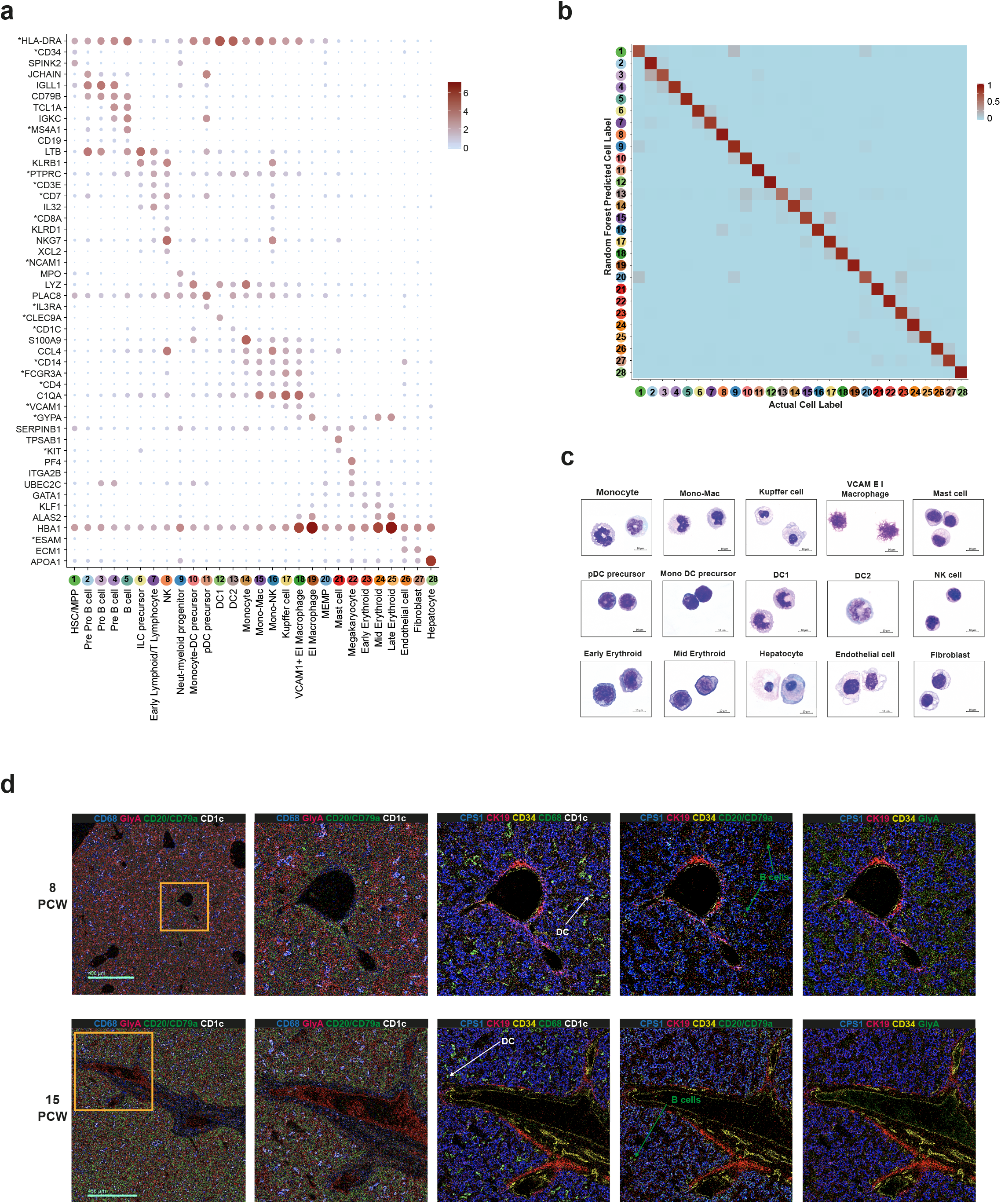
Multi-modal and spatial validation of cell types. **a**, Dot plot of selected 48 marker genes for each liver cell type from **1c**; * indicates markers used for FACS-isolation of cells. Gene expression frequency indicated by spot size and level by colour intensity. **b**, Assessment of fetal liver cell markers from **a** using random forest to assign cell labels, actual = using all genes and predicted = using selected 48 genes. **c**, Giemsa stained cytospin prepared cells isolated by FACS based on markers with * in **a** (images representative from n=3). **d**, Overlaid, pseudo-coloured Hyperion images for 8 PCW and 15 PCW fetal liver. Far left images are shown at 5x magnification with zoom of insets on right at 20x magnification (1 μm/pixel).

Next, we devised a panel of markers to evaluate the spatial distribution of the erythroid, mast cell, myeloid and lymphoid lineages within the same fetal liver tissue section using imaging mass cytometry (Figure 2d). Islands of haematopoietic cells are interspersed in the sinusoids around hepatocyte aggregates throughout the fetal liver. CD68^+^ macrophages within the sinusoids were surrounded by GlycophorinA^+^ erythroid cells (Figure 2d). CD1c^+^ DCs and CD79a/CD20^+^ cells from the B-lineage are sparsely distributed (Figure 2d). The liver architecture evolved considerably between 8 and 15 PCW. Loose aggregates of hepatocytes in early 8 PCW fetal liver are more densely organised at 15 PCW but hepatic lobules with a distinct central vein and portal triad are not clearly observed. Bile duct development becomes evident between 8 and 15 weeks (Figure 2d). Although the relative proportion of haematopoietic cells approximated our scRNA-seq profile, there were more hepatocytes observed by imaging mass cytometry (Figure 2d and Figure 1c). This is in keeping with published data on the fragility of hepatocytes following *ex vivo* isolation and high expression of mitochondrial genes (up to 50%) in scRNA-seq analysis of adult hepatocytes^16–18^. By validating our single cell transcriptome datasets with multiple modalities including flow cytometry, morphology and imaging mass cytometry for spatial resolution, we provide an integrated map of haematopoietic cells in the fetal liver.

### Fetal liver and NLT haematopoiesis

Following *ab initio* identification of fetal liver cells, we leveraged the real-time data from liver samples across the gestational stages to infer developmental trajectories of fetal liver haematopoietic cells. For all lineages, we used three different computational approaches (force-directed graph, diffusion map and approximate graph abstraction (AGA) using Scanpy) and derived differentially expressed genes over pseudotime using Monocle. Force-directed graph visualisation guided us to identify three distinct connections to a central HSC/MPP node (Figure 3a and Supplementary Video 1). These connections correspond to cells from the following lineages; the erythroid-megakaryocyte-mast cell, B-cell and innate/T-lymphoid, and myeloid lineages, and provide a comprehensive view of putative haematopoietic differentiation trajectories in human fetal liver (Figure 3a). The central location of HSC/MPPs and distribution of cells from the respective lineages in Figure 3a support the validity of our cell annotation (Figure 1c).

**Figure 3:**
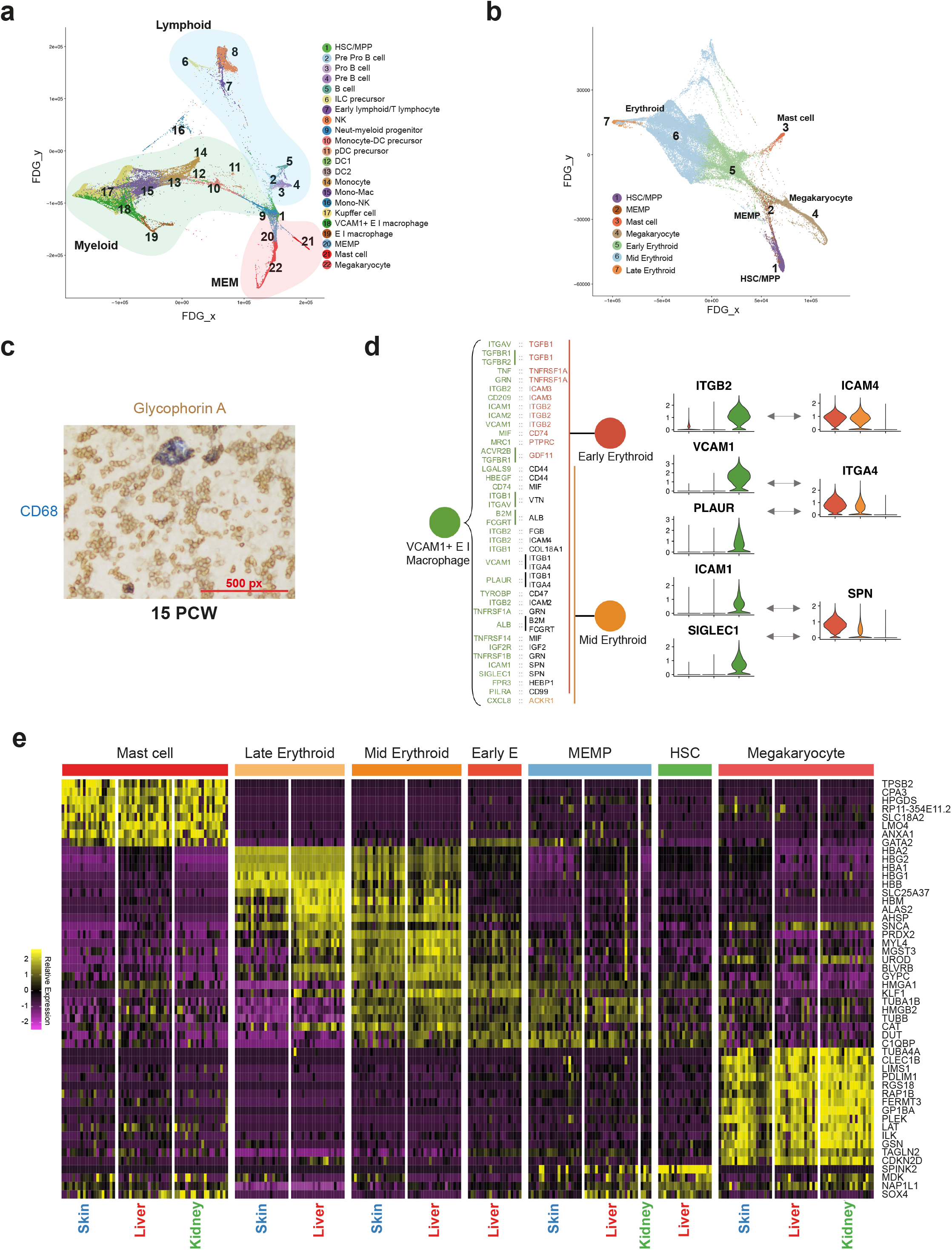
Fetal liver and NLT haematopoiesis. **a**, Force-directed graph (FDG) visualisation of all haematopoietic cell from **1c**. **b**, FDG visualisation of fetal liver HSC/MPP, erythroid, megakaryocyte and mast cell lineages from **a**. **c**, Immunohistochemical staining with GlycophorinA and CD68 of fetal liver (representative of n=3). **d,** Receptor-ligand interactions from CellPhoneDB between VCAM1^+^ EI macrophages and two erythroid populations (early and mid) with violin plots showing gene expression value of each receptor and ligand. Violin plot colours correspond to the predicted interacting cell types. **e**, Heat map of the top 20 expressed genes in fetal liver and NLT subsets (max 20 cells per subset), relative expression values from low (purple) to high (yellow).

We next focused on each of the three haematopoietic branches radiating from the HSC/MPP node in fetal liver and interrogated their relationship to haematopoietic cells in fetal NLT (skin and kidney) (Figure 3a and Supplementary Figure 3b). For comparison between fetal liver and NLT haematopoietic cells, we used AGA scores to evaluate similarities across tissues and assessed the differentially expressed genes between tissue counterparts (see Methods). In the fetal liver, all three trajectory inference approaches reveal a consistent result, inferring mast cells, erythroid and megakaryocyte lineages origin from a shared progenitor downstream of HSC/MPP, the putative megakaryocyte-erythroid-mast cell progenitor (MEMP) (Figure 3b and Extended Data Figure 3a). The unexpected observation of mast cell coupling with erythroid fate is in keeping with observations of mast cell and basophil origin during murine bone marrow and human cord blood haematopoiesis^19,20^. However, yolk sac-derived mast cells have also been observed in murine skin and connective tissues suggesting potential alternative origin for mast cells in NLT^19–22^.

We can discern three transcriptionally distinct stages of erythroid development across a continuum characterised by early, mid and late erythroblasts, with variable expression of haemoglobin genes (Figure 1c and Extended Data Figure 3b). As fetal liver erythropoiesis is known to be supported by adhesive interactions within erythroblastic islands (Figure 3c)^23^, we used a quantitative statistical method (cellphonedb.org and Vento-Tormo et al., Nature *In press*) to predict cell surface receptor-ligand interactions between erythroblasts and erythroblastic island (EI) macrophages. Our analysis identified molecules previously implicated in haematopoiesis such as *VCAM1, ITGB1, ITGB2, ITGA4*, *SIGLEC1, ICAM4* and *SPN* (Figure 3d)^23–28.^ With our high-resolution data, we can now pinpoint these molecules as supporting the differentiation of erythroblasts by EI macrophages (Figure 3d).

We next evaluated the genes that were differentially regulated during the transition of HSC/MPPs into erythroid cells, megakaryocytes and mast cells (Extended Data Figure 3c). Genes dynamically expressed during erythroid differentiation include the known transcription factors (TFs) *TAL1* and *GATA2*; *BPGM*, which binds haemoglobin and the glycolysis enzyme *PKLR* (Extended Data Figure 3c)^29,30^.

In contrast, megakaryocyte differentiation involves temporal regulation of *F2R*, *PBX1* and *MEIS1*, the latter known to dimerise and activate platelet factor 4 (PF4) (Extended Data Figure 3c)^31^. Interestingly, we show that *HES1*, previously reported to drive mast cell differentiation through Notch signalling is dynamically regulated during fetal liver mast cell differentiation (Extended Data Figure 3c)^32^. Considering our data from the NLTs skin and kidney, we show that mast cells, megakaryocytes, putative MEMP, and erythroid cells are also present in NLT, but HSC/MPPs are absent (Supplementary Figure 3 and Extended Data Figure 3d-e). Since HSC/MPPs are confined to fetal liver, the presence of MEMP, megakaryocytes and mast cells in all 3 sites with similar single cell transcriptomes suggests that these cells may circulate between tissues during this stage of fetal life. Megakaryocytes are more abundant in fetal kidney, but early, mid and late erythroid cells are only present in skin (Extended Data Figure 3d-e).

We compared highly expressed and differentially expressed genes of corresponding cell types in fetal liver, skin and kidney (Figure 3e). The transcriptome of mast cells is virtually identical across all three tissues, as is the transcriptome of megakaryocytes, consistent with potential migration of these cells between sites (Figure 3e).

The similarity in differentiation trajectories between cells in the two NLT sites and the fetal liver implies one of two possibilities: either (i) there is local maturation of progenitors in NLT or (ii) cells at several differentiation stages exit from the fetal liver and independently migrate to the NLT (Extended Data Figure 3e). The absence of erythroid cells downstream of putative MEMP in fetal kidney suggests that extra-hepatic erythropoiesis is tissue-restricted (Figure 3e and Extended Data Figure 3e).

To further evaluate if putative MEMP may have expansion capacity in NLT, we inspected the expression levels of the proliferation (*MKI67)* and cell cycle genes (Seurat CellCycleScoring, see Methods) (Extended Data Figure 3f). Amongst putative MEMP, 50-70% are in S and G2/M phase in fetal skin and kidney, while only 25% are in S and G2/M phase in fetal liver (Extended Data Figure 3f). Furthermore, in contrast to fetal liver and kidney, putative MEMP in skin also express some early erythroblast genes including *MYL4* (Figure 3e)^33,34^, suggesting that these may act as progenitors *in situ* in the skin. These findings imply that during early development, the skin in physiological state can contribute to erythropoiesis and supplement fetal liver erythroid output.

### Lymphoid lineages in fetal liver and NLT

Previous studies have reported the presence of T and B lymphocytes^35^, NK cells^36^, and ILCs^37^ in the human fetal liver. We observe two lymphoid branches which include cells from the B-lineage and NK/T/ILC lineage respectively (Figure 4a and Extended Data Figure 4a). Interestingly, the ‘early lymphoid/T lymphocyte’ cluster comprise heterogeneous cells which vary by gestation stage (Extended Data Figure 1c, Figure 4a and Extended Data 4b-c).

**Figure 4:**
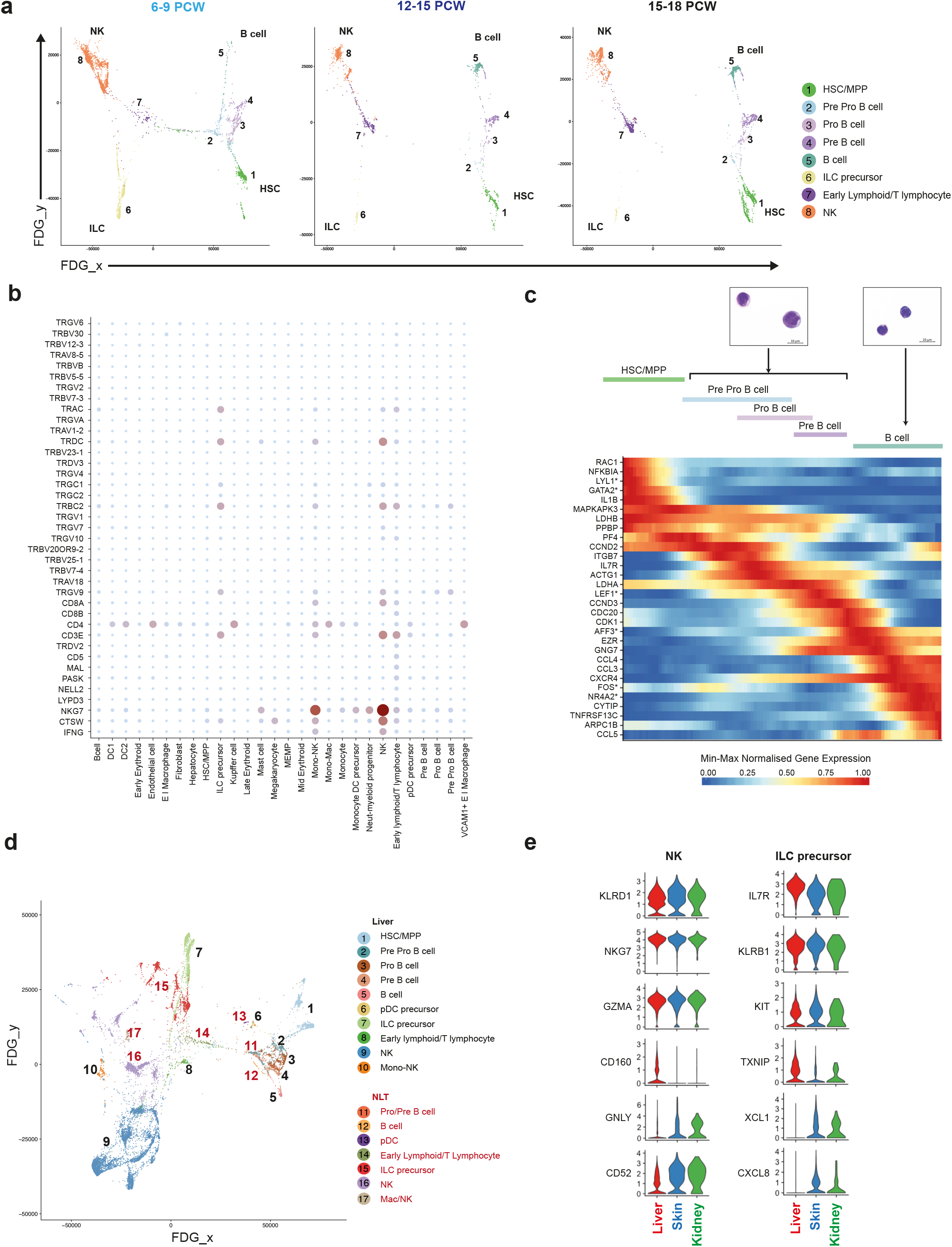
Lymphoid lineages in fetal liver and NLT. **a**, FDG visualisations of fetal liver HSC/MPP and lymphoid cell types from **1c** showing changes over developmental stages (6-9 PCW, 12-15 PCW and 15-18 PCW). **b**, Dot plot showing expression of V(D)J transcripts in fetal liver cell types (n=6). Gene expression frequency indicated by spot size and level by colour intensity. **c**, Dynamically expressed pseudotime genes during B cell development, transcription factors are marked with *. Giemsa stained Pro/Pre B cells and B cells prepared by cytospin from **2c** (representative of n=3). **d**, FDG visualisation of F1-F10 fetal liver and corresponding skin and kidney lymphoid cells. **e**, Violin plots showing expression of shared and differentially expressed genes in NK and ILC precursors across F1-F10 fetal liver, skin and kidney.

Between 6-9 PCW, before T cells emerge from the thymus, the cells within this cluster express *GATA3*, *KLRB1*, *CD3D, CD3E, CD7* and *JCHAIN* (Extended Data Figure 4b-c), and may relate to the previously reported putative fetal liver thymocyte progenitors, which are capable of generating αβT cells upon co-culture with thymic epithelial cells^38–40^. After 12 PCW, the cells in this cluster are characterised by the expression of *IL7R, CD2, CD7, CD27*, *CD8B* with some cells co-expressing *CD4* (Extended Data 4b-c). Lymphocyte antigen receptor gene analysis (from the 10x Genomics 5’ V(D)J kit) shows T cell receptor delta constant chain (*TRDC*) and T cell receptor alpha constant chain (*TRAC*) expression by cells from 12-14 PCW but cells from 14-18 PCW lack *TRDC* expression (Figure 4b and Extended Data Figure 4c).

The *TRDC/TRAC* expressing cells lack *GZMB* and *PRF1*, the cytoplasmic granular products characteristic of mature CD8^+^ T cells (Figure 4b and Extended Data 4c). These findings are in keeping with the seeding of fetal liver by γδT cells and αβT cells sequentially following their exit from thymus after 12 PCW^41^. This is consistent with previous reports of T cell identification only after 18 PCW^40,42^.

Using diffusion map and AGA analyses, we show that NK cells (characterised by *NCAM1*, *EOMES, CD7*, *IL2R* and *CD3E*) and ILC precursors (characterised by *KIT*, *KLRB1*, *IL7R*, *ID2, AHR, RORC* but lack *IL2R*, *TBX21*, *IL1RL1* (*ST2*)) are connected *via* the lymphoid branch (Figure 2a, 4a and Extended Data Figure 4a). This finding is in keeping with existing literature of a shared progenitor for NK and ILCs in human and mouse^43–45^. Intriguingly, we observe a cluster of cells which express the gene modules of monocytes and NK cells (Mono-NK cluster), similar in characteristics to a monocyte subset previously identified in adult human peripheral blood^46^.

B-lineage cells across a continuum of differentiation states are present in the fetal liver (Figure 1c, Figure 3a and 4a). The most primitive cluster expressing *CD34*, *SPINK2* and B-lineage transcripts *IGLL1* and *PAX5* is designated ‘pre pro-B’. The ‘pro-B’ and ‘pre-B’ cell clusters have sequentially higher expression of B cell related transcripts *EBFI*, *MS4A1*, *CD79B, DNTT* and MHC-II genes such as *HLA-DRA,* while expression of *JCHAIN* and *LTB* declines (Figure 2a). The primitive state of ‘pro/pre-B’ clusters is confirmed by their high nuclear to cytoplasmic ratio, immature chromatin and presence of nucleoli (Figure 4c). Genes involved in the transition from HSC/MPPs to B cells include *RAC1*, *JUN* and *LDHB* (Figure 4c).

Analysis of the lymphoid lineage across gestational age reveals the presence of pre-B cells between 6-9 PCW, but their differentiation into mature B cells in the fetal liver is efficiently completed only after 9 PCW (Figure 1e and 4a). We noted *IL10RB-IL10* receptor-ligand expression, known to be important for B cell differentiation^47^, only after 9 PCW in fetal liver. The larger question of cell-intrinsic *versus* tissue-microenvironment factors relating to gestational age support for B cell differentiation requires further investigations.

We next evaluated the composition and molecular characteristics of lymphoid cells in fetal NLT and evaluated transcriptionally similar cell types to fetal liver lymphoid cells, which yield high AGA connectivity scores (Figure 4d and Extended Data Figure 4d). Cells resembling Mono-NK are present in the skin but have greater expression of macrophage genes *LYVE1*, *F13A1, C1QA/B/C* and *RNASE1* than their fetal liver counterparts (Extended Data Figure 4e). Pro-B, pre-B and B cells are present in NLT but HSC/MPPs and pre pro-B cells are absent (Figure 4d).

NK precursors, NK cells and ILC precursors present in NLT share a transcriptional signature with their liver counterparts, however tissue-specific expression of chemokine (*XCLI, CXCL8*) and cytotoxic granule genes (*GNLY*) suggests maturation and tissue adaptation in the skin and kidney (Figure 4e). ILC precursors in NLT lack the full characteristic markers and TFs of their mature progenies; ILC1, ILC2 and ILC3 (Extended Data Figure 4f). Together, these findings suggest that NLTs are seeded by NK and ILC precursors from fetal liver, which differentiate *in situ* and acquire tissue-related gene expression profile.

### Tissue signatures in developing myeloid cells

In mice, elegant fate-mapping studies have demonstrated that tissue macrophages are seeded by yolk sac and fetal liver progenitors^48,49^. In contrast, dendritic cells (DCs) are thought to originate from bone marrow-derived HSC/MPPs through a monocyte-independent lineage^50,51^. Our data show the presence of myeloid progenitors, monocytes, DCs and macrophages as early as 6 PCW in the fetal liver and also in NLT, revealing the ability of fetal liver HSC/MPPs to also generate DCs (Figure 1c, Extended Data Figure 1c and Supplementary Figure 3b). It is unclear if fetal liver HSC/MPP-derived DCs are homeostatically and functionally distinct from BM-HSC/MPP-derived DCs.

Diffusion map and AGA analysis of fetal liver myeloid cells reveal transcriptionally related cells stemming from HSC/MPP with increasing acquisition of myeloid genes (Figure 5a and b and Extended Data Figure 5a). The earliest neutrophil myeloid progenitor expresses *CD34*, *SPINK2*, and known myeloid markers including *GATA2*, *ELANE*, *MPO* and *LYZ* (Figure 2a). Monocyte-DC precursors share expression of monocytes and DC genes *S100A9*, *CD1C* and *LYZ* but do not express *CD14*. Monocytes express *CD14*, *S100A9*, *CD68* but not *CD1C*. Conventional DCs express the canonical markers for the respective subsets; *CD1C* in DC2 and *CLEC9A*, *CADM1* and *XCR1* in DC1 (Figure 2a).

**Figure 5:**
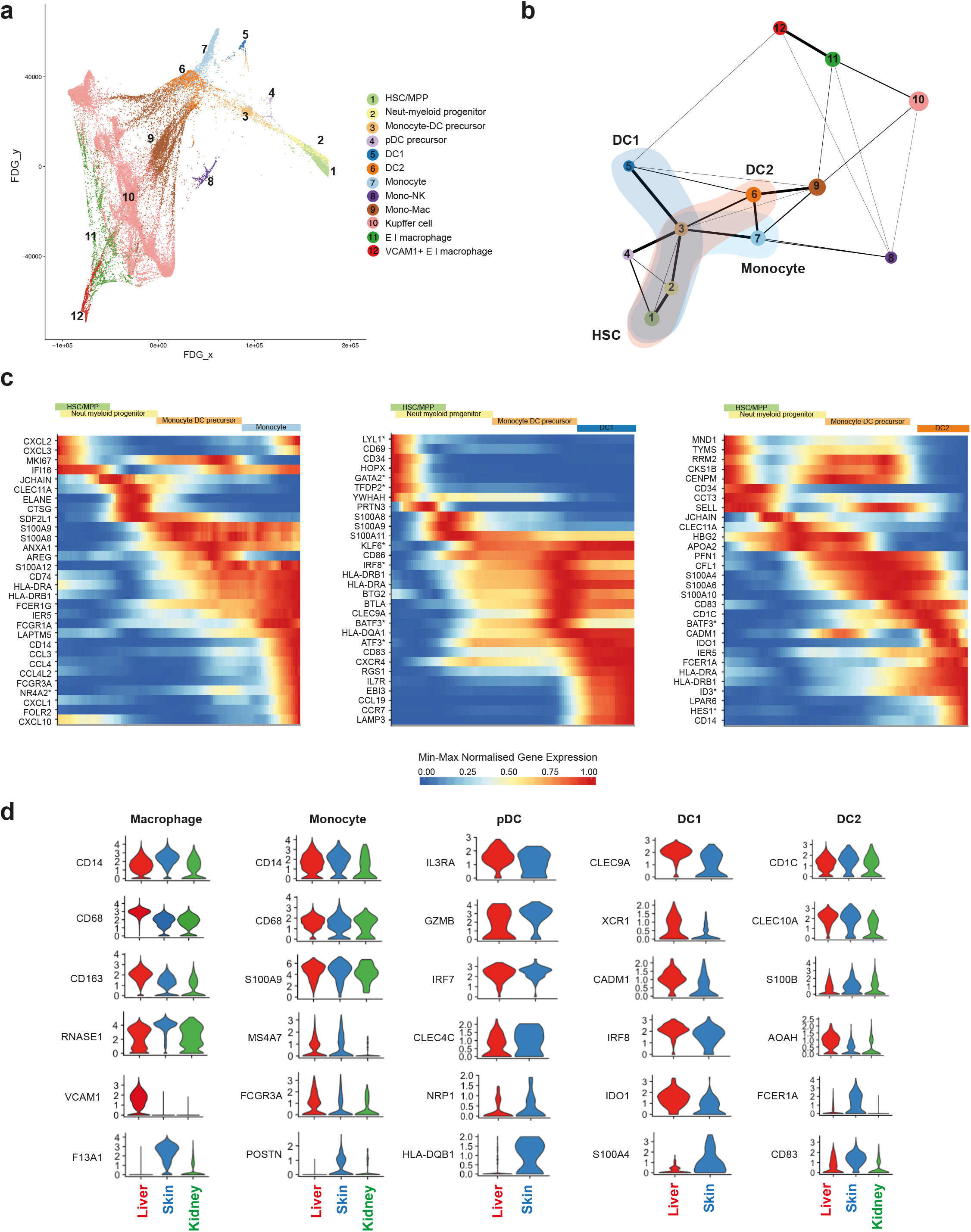
Tissue signatures in developing myeloid cells. **a**, FDG visualisation and **b,** AGA analysis of fetal liver HSC/MPP and myeloid cells from **1c**. Colour clouds highlight DC1, DC2 and monocyte lineages depicted in **Figure 5c**. **c**, Dynamically expressed pseudotime genes during monocyte, DC1 and DC2 development, transcription factors marked with *. **d**, Violin plots showing expression values of shared and differentially expressed genes in corresponding myeloid populations across fetal tissues (F1-F10).

Diffusion map and AGA analysis suggests the bifurcation of DC1, DC2 and monocytes from monocyte-DC precursors (Figure 5a and 5b, Extended Figure 5a). However, pDCs appear to branch off from two cell clusters: HSC/MPP-early pro-B cells and monocyte-DC precursors (Figure 5a and Extended Data Figure 5a). This is in keeping with recent reports in mice of their mixed lymphoid and myeloid origin^19,52–54^.

AGA analysis shows only weak edges (connecting lines where thickness quantitates transcriptional similarities) between fetal liver macrophages (Kupffer cells and EI macrophages) and monocyte-DC precursors (Figure 5b). This indicates that fetal liver HSC/MPPs are the likely source of DCs, monocytes and mono-mac cells, but that Kupffer cells and EI macrophages may arise from an alternative source such as yolk sac-derived progenitors. Yolk sac progenitors have been shown to contribute to tissue macrophages, including Kupffer cells, in prenatal mice^55^ which can survive until adulthood depending on the tissue and subsequent niche availability^56–58^.

We next evaluated the genes dynamically expressed during the transitions from HSC/MPPs to DCs and HSC/MPPs to monocytes-macrophages independently (Figure 5c). TFs previously implicated in DC differentiation such as *GATA2*, *IRF8* and *BATF3*^51^ are dynamically regulated in pseudotime. We infer a role for additional transcription factors in these lineages: *ATF3*, *HES1* and *ID3*. In addition to canonical genes such as *CD14* and *FCGR1A/3A*, we highlight a role for the previously unappreciated nuclear receptor family member *NR4A2* during monocyte development.

In NLTs, monocytes, macrophages and DC2 are present in both skin and kidney, pDC and DC1 are only detected in skin but we cannot exclude their presence at low frequency in the kidney (Supplementary Figure 3b). Corresponding clusters in NLT and fetal liver shared transcriptional similarity (Extended Data Figure 5b). Tissue-specific differences in gene expression were most notable in macrophages: there is unique expression of *VCAM1* and *F13A1* in liver^26^ and skin^59^ macrophages respectively, resembling the profile of adult macrophages in these respective tissues (Figure 5d).

NLT pDCs and cDCs express DC maturation and activation markers compared to fetal liver DCs (Figure 5d and Extended Data Figure 5c). Skin pDCs had a higher expression of MHC-II genes, *CLEC4C* (CD303) and *NRP1* (CD304) than fetal liver pDCs, suggesting a more mature state (Figure 5d and Extended Data Figure 5c). DC1 in skin express more *S100A4,* a molecule involved in T cell activation, than liver DC1. The expression of *AOAH*, an enzyme involved in lipopolysaccharide response modulation, was more specific to liver DC2, while skin DC2 expressed more *FCER1A* and *CD83* (Figure 5d). These tissue-related gene signatures suggest acquisition of functional specialisations by macrophages and DCs during early development. The appearance of activated DCs despite the sterile fetal environment suggest an active role for fetal DCs in mediating tolerance as previously reported^60^.

### HSC/MPP differentiation potential by gestation

Our observation of a central HSC/MPP cluster from which the earliest lineage-committed cells radiate is in keeping with recent observations from scRNA-seq analysis in post-natal mice and humans^19,61,62^ (Figure 6a, Extended Data Figure 6a-c). Genes dynamically regulated during HSC/MPP transition to the early progenitors include recognised TFs and receptors implicated during haematopoiesis such as *ELANE*, *LYZ*, *MPO* and *AZU1* for neutrophil-myeloid lineage, *LTB*, *JCHAIN*, *IL7R* and *VPERB1* for the lymphoid lineage, and *GATA1*, *GATA2*, *EPOR* and *FCER1* for megakaryocyte-erythroid-mast cell lineage (Figure 6b).

**Figure 6:**
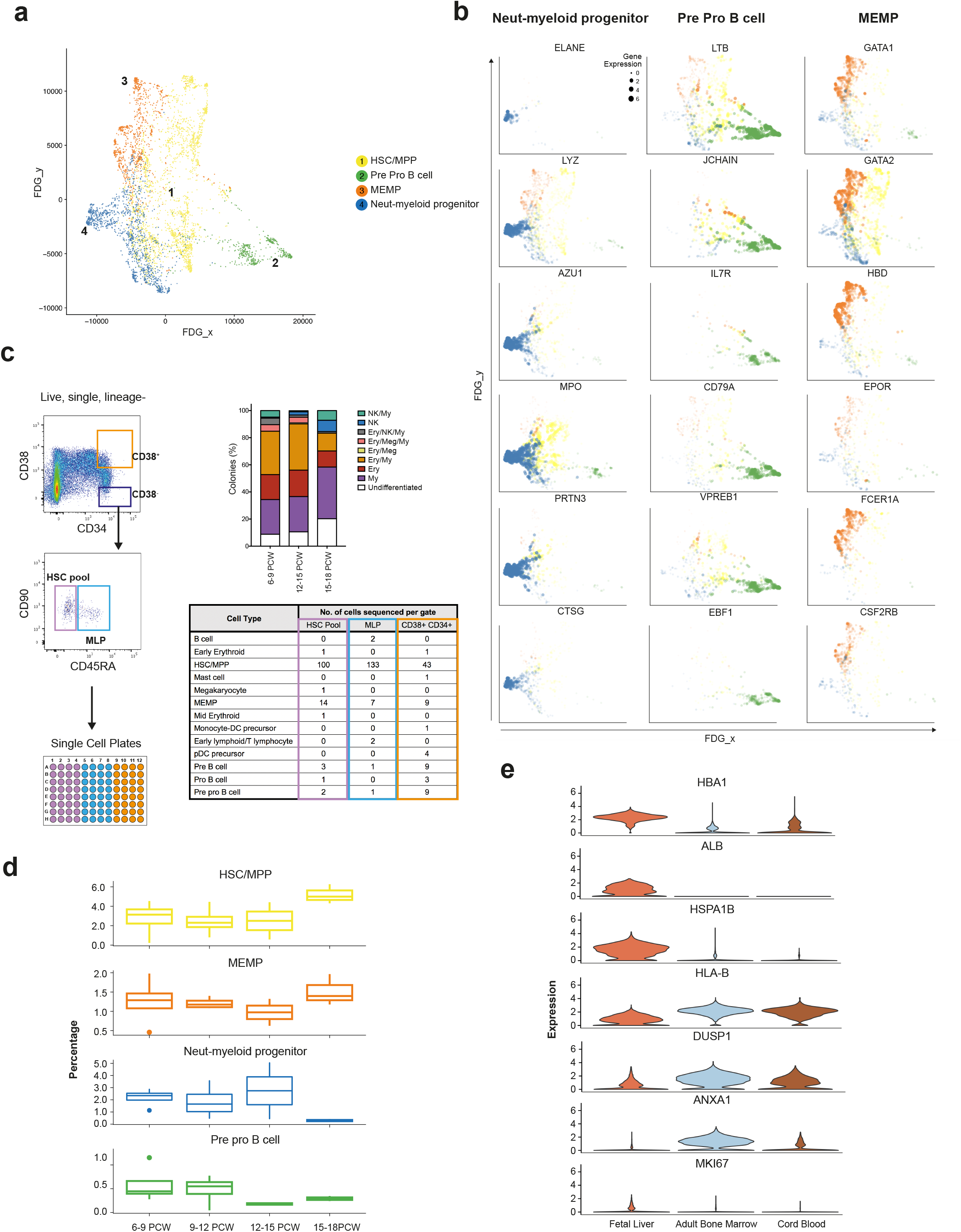
HSC/MPP differentiation potential by gestation. **a**, FDG visualisation of HSC/MPP and early haematopoietic progenitor populations from **1c**. **b**, Feature plots of dynamically expressed pseudotime genes during HSC/MPP transition to neutrophil-myeloid, MEMP and pre pro B cells. Size of dots indicate gene expression level (small = low; large = high). **c**, Left panel shows experimental design for single cell transcriptome and culture of fetal liver cells from FACS gates illustrated, top right shows % colonies cultured from ‘HSC pool’ gate; 96 cells cultured from each sample (n=5), bottom right panel shows scRNA-seq-based cell identities for 349 cells from FACS gates corresponding to table column colour (n=3). **d,** Percentage (median +/− SD) of HSC, neutrophil-myeloid progenitors, MEMP and pre pro B cells in fetal liver during development (n=13). **e,** Violin plots (expression values) of differentially expressed genes between HSC/MPPs in fetal liver (n=13), cord blood (n=8) and adult bone marrow (n=8).

We hypothesised that the cellular composition of the developing fetal liver resulted from local modulation of HSC/MPP potential. To test this hypothesis, we FACS-isolated single cells from the CD34^+^CD38^+^, CD34^+^CD38^−^CD90^+^CD45RA^−^ and CD34^+^CD38^−^CD90^+^CD45^+^ FACS gates and profiled them by both plate-based single cell transcriptomics (Smart-seq2 protocol) and single cell clonal differentiation potential analysis using established culture systems^63^ (Figure 6c).

We used an SVM trained on the entire fetal liver dataset to identify HSC/MPPs and lineage-committed progenitors captured by Smart-seq2 analysis. By aligning transcriptome profiles with FACS index metadata used to isolate single cells (Figure 6c), we confirm the enrichment of HSC/MPPs (~85%) within the CD34^+^CD38^−^ gate further supporting our annotation of fetal liver cells (Figure 6c and Extended Data Figure 6d). The majority of cells within the CD34^+^CD38^−^CD90^−^CD45RA^+^ MLP gate also contained HSC/MPP, in agreement with previous reports of human cord blood MLP being transcriptionally closest to HSC/MPP^64^.

Differentiated progenies from the single cell culture were also analysed by flow cytometry to establish the differentiation potential of each single cell (Extended Data Figure 6e and Supplementary Figure 1c). Single cells cultured from the CD34^+^CD38^−^ gate (HSC/MPP enriched) exhibited mutipotential differentiation capacity (Extended Data Figure 6e). The proportion of colonies derived from CD34^+^CD38^−^CD45RA^−^ single cells containing NK cells significantly increased with gestational age, whilst the proportion of colonies containing erythroid/megakaryocytic cells significantly decreased (Extended Data Figure 6f).

We propose this differentiation skew as an HSC/MPP-intrinsic property, since liquid culture conditions without stroma produced the same ratios in output (Extended Data Figure 6g). We also observed a trend towards reduced multipotentiality with increased gestational age (Extended Data Figure 6e). These findings support the hypothesis of differential HSC/MPP intrinsic potential by gestational stage and is in keeping with our earlier observation of the relative decrease in erythroid cells during development (Figure 1d).

We observed a relative increase in the proportion of neutrophil-myeloid progenitors and decline in pre pro-B cells up to 14 PCW (Figure 6d) followed by the rapid decline of the former between 14-18 PCW. This was in contrast to a rapid increase in the relative frequency of HSC/MPPs >14 PCW. The relative decline in early progenitors and expansion in HSC/MPP >14 PCW may be due to proliferative phase in preparation for HSC/MPP seeding the bone marrow niche or the exit of the early progenitors (neutrophil-myeloid, pre pro-B) to BM or even NLT^65–67^. Comparative analysis of fetal liver and fetal bone marrow in future studies may provide new insights into the relationship of HSC/MPPs and early progenitors across these tissues. Differential expressed genes analysis between HSC/MPPs in fetal liver, CB and adult BM reveals higher expression of *MKI67 i*n fetal liver HSC/MPPs suggestive of enhanced proliferative potential as previously reported in mice^65–67^, increased expression of liver-specific genes (*ALB*), propensity for erythropoiesis (*HBA1)* and heat shock protein (*HSPA1B*) potentially conferring tissue identity and maintenance of genome and proteome integrity (Figure 6e and Extended Data Figure 6h). FL HSC/MPPs expressed lower levels of MHC-I (*HLA-B*) suggesting reduced antigen presenting potential compared to CB and adult BM HSC/MPPs (Figure 6e).

Collectively, our findings demonstrate that intrinsic changes in HSC/MPP numbers and differentiation potential occur over the first and second fetal trimesters. These changes are likely to be pivotal in allowing fetal liver haematopoiesis to adapt to the needs of the developing fetus; first the establishment of an effective oxygen transport system and subsequently the development of a complete blood and immune system.

## Discussion

The development of the human immune system *in utero* has remained poorly understood. Previous studies have generally focused on a single immune cell type or lineage, and deployed markers of adult immune cells to identify fetal immune cells. Using single cell transcriptome profiling, we were able to overcome prior assumptions that adult markers faithfully identify fetal cells and resolve the cellular heterogeneity within human fetal liver. Analysis of fetal liver, skin and kidney over gestation enabled us to abstract dynamic temporal information across tissues to comprehensively establish the haematopoietic and immune cell composition during fetal liver haematopoiesis.

We demonstrate the power and validity of transcriptome-based cell classification by integrating the data with multiple modalities: FACS, Giemsa staining for morphology, and imaging mass cytometry-based multiplex protein measurements. These methods allow for cost-effective and scalable isolation of cell types, and analysis of their spatial localisation.

Our approach highlights key insights; intimate linking of mast cell origin with erythro-megakaryopoiesis, erythropoiesis in fetal skin as a normal feature rather than stress-response to disease, establishment of DC network as early as 6 PCW, potential dual myeloid and lymphoid origin of pDCs and tissue adaptation of NKs, ILCs, DCs and macrophages during development. Our findings reveal modulation of HSC/MPP intrinsic differentiation potential over gestation age as a functional mechanism to regulate haematopoietic output of the fetal liver throughout the first and second trimesters.

While the fetal liver represents the primary site of fetal erythropoiesis, our data suggest that fetal skin also contain proliferating erythroid progenitors which are likely to contribute to the high demand for blood development in the fetus. This is supported by the presence of skin lesions containing extramedullary haematopoiesis in fetuses and neonates with severe fetal anaemia^68^. Although the percentage of erythroid lineage cells in skin is 1% of that in fetal liver, the total skin surface area of a fetus would support a significant contribution to erythropoiesis particularly in the setting of increased erythropoietic need. Extramedullary haematopoiesis in the skin may also occur in congenital leukaemias, congenital CMV infection and in adults with myelofibrosis where megakaryocyte-containing papules are observed^69^. Circulating primitive erythroblasts have been observed in the mouse^70^ and significant platelet biogenesis in the post-natal murine lung has also been previously reported^71^.

The presence of MEMP, megakaryocyte and erythroid cells at different maturation stages in 6-12 PCW NLT suggest local maturation of progenitors, or extrahepatic haematopoiesis. The acquisition of tissue-specific molecular features is well recognised for myeloid and lymphoid cells including ILCs and T cells^49,72,73^. This may relate to functional adaptation of myeloid and lymphoid cells but a more generic function across tissues for mast cells, which are transcriptionally unaffected by their local tissue microenvironment.

In summary, our comprehensive atlas of human fetal liver blood and immune cells provides a foundational resource for understanding fetal liver haematopoiesis and the developing immune system. Our reference dataset will be invaluable for studies on paediatric blood and immune disorders and exploiting HSC/MPPs for therapy. Our approach using single cell transcriptomics to study human development provides a framework that can be applied to study any temporal processes across the human lifespan.

## Materials and methods

### Tissue Acquisition

Human fetal tissues were obtained from the MRC/Wellcome Trust-funded Human Developmental Biology Resource (HDBR; http://www.hdbr.org)^74^ with appropriate written consent and approval from the Newcastle and North Tyneside NHS Health Authority Joint Ethics Committee (08/H0906/21+5). HDBR is regulated by the UK Human Tissue Authority (HTA;www.hta.gov.uk) and operates in accordance with the relevant HTA Codes of Practice.

### Tissue Processing

All tissues were processed immediately after isolation using the same protocol. Tissue was transferred to a sterile 10mm^2^ tissue culture dish and cut into <1mm^3^ segments before being transferred to a 50mL conical tube. Tissue was digested with 1.6mg/mL collagenase type IV (Worthington) in RPMI (Sigma-Aldrich) supplemented with 10%(v/v) heat-inactivated fetal bovine serum (Gibco), 100U/mL penicillin (Sigma-Aldrich), 0.1mg/mL streptomycin (Sigma-Aldrich), and 2mM L-Glutamine (Sigma-Aldrich) for 30 minutes at 37°C with intermittent shaking. Digested tissue was passed through a 100μm filter, and cells collected by centrifugation (500g for 5 minutes at 4°C). Cells were treated with 1X RBC lysis buffer (eBioscience) for 5 minutes at room temperature and washed once with flow buffer (PBS containing 5%(v/v) FBS and 2mM EDTA) prior to counting.

### Fetal developmental stage assignment and chromosomal assessment

Embryos up to 8 post conception weeks (PCW) were staged using the Carnegie staging method^75^. After 8 PCW, developmental age was estimated from measurements of foot length and heel to knee length and compared against a standard growth chart^76^. A piece of skin, or where this was not possible, chorionic villi tissue was collected from every sample for Quantitative Fluorescence-Polymerase Chain Reaction analysis using markers for the sex chromosomes and the following autosomes 13, 15, 16, 18, 21, 22, which are the most commonly seen chromosomal abnormalities.

### Flow cytometry and FACS for scRNA-seq

Antibody panels were designed to allow enrichment of cell fractions for sequencing and cell types validation. Antibodies used for FACS isolation are listed in Supplementary Table 1. An antibody cocktail was prepared fresh by adding 3μL of each antibody in 50μL Brilliant Stain Buffer (BD) per tissue. Cells (≤10×10^6^) were resuspended in 50-100μL flow buffer and an equal volume of antibody mix was added to cells from each tissue. Cells were stained for 30 minutes on ice, washed with flow buffer and resuspended at 10×10^6^cells/mL. DAPI (Sigma-Aldrich) was added to a final concentration of 3μM immediately prior to sorting. Flow sorting was performed on a BD FACSAria™ Fusion instrument using DIVAv8, and data analysed using FlowJoV10.4.1. Cells were gated to remove dead cells and doublets, and then isolated for scRNA-seq analysis (10x or Smart-seq2). For 10x, cells were sorted into chilled FACS tubes coated with FBS and prefilled with 500μL sterile PBS. For Smart-seq2, single cells were index-sorted into 96-well lo-bind plates (Eppendorf) containing 10μL lysis buffer (TCL (Qiagen) + 1% (v/v)B-mercaptoethanol) per well.

B cells were also investigated by flow cytometry as per Roy et. al.,^77^.

### Cytospins and mini bulk RNA-seq validation

Frozen suspended cells from human fetal livers (F15 and F18) were thawed, counted and split for staining with two separate panels (see Supplementary Table 2 for antibody details). Cells were stained for 30 minutes on ice followed by DAPI staining. FACS was performed on a BD FACSAria™ Fusion instrument using DIVAv8, and data analysed using FlowJov10.4.1. Cells were isolated into chilled FACS tubes coated with FBS and prefilled with 500μL sterile PBS for cytospin (500 – 2000 cells), or into 1.5mL microfuge tubes containing 20 μL lysis buffer (100 cells). Giemsa staining (Sigma-Aldrich) was used to determine morphology of sorted cells on cytospins. Slides were viewed using a Zeiss AxioImager microscope, images taken of 4 fields from n=3 samples using the 100x objective, and viewed using Zen (v2.3).

### HSC/MPP Culture

MS5 in log-phase growth (DSMZ, Germany, passage 6-10) were seeded into 96-well flat-bottom plates (Nunclon delta surface; Thermo) at a density of 3000 cells per well 24 hours prior to sorting. Medium was Myelocult H5100 (Stem Cell Technologies) supplemented with 100U/mL Penicillin and 0.1mg/mL Streptomycin (Sigma-Aldrich). On the day of sorting, media were replaced with Stem Pro-34 SFM media (Life Technologies) supplemented with 100U/mL Penicillin and 0.1mg/mL Streptomycin, 2mM L-glutamine (Sigma-Aldrich), stem cell factor 100ng/ml (Miltenyi), Flt3 20ng/ml (Miltenyi), TPO 100ng/ml (Miltenyi), EPO 3ng/ml (Eprex), IL-6 50ng/ml (Miltenyi), IL-3 10ng/ml (Miltenyi), IL-11 50ng/ml (Miltenyi), GM-CSF 20ng/ml (Miltenyi), IL-2 10ng/ml (Miltenyi), IL-7 20ng/ml (Miltenyi) and Lipids (hLDL) 50ng/ml (Life Technologies)^63^.

Three populations of HSC/MPPs and progenitors were isolated from fetal liver suspension (1 fresh F15, 2 frozen F18, F19). Populations were identified from the DAPI^−^, doublet-excluded gate as CD3/CD16/CD11c/CD14/CD19/CD56^−^, CD34^+^ cells (see Supplementary Table 3 for antibody details). The HSC/MPP pool and MLP were found within the 20% of cells with lowest CD38 expression: HSC/MPP pool were CD90^+/−^ and CD45RA^−^ whilst MLP were CD90^−^CD45RA^+^. Progenitors with the highest 20% of CD38 expression were sorted for comparison. Single cells were sorted using a BD FACSAria™ Fusion operating DIVAv8, and sorted directly onto MS5 for culture, or into 96-well lo-bind plates containing 10 μl/well lysis buffer for Smart-seq2 scRNAseq. Single-cell-derived colonies analysis was performed as described by the Laurenti Lab^63^. In brief, colonies were harvested into 96 U-bottom plates using a plate filter to prevent the carryover of MS5 cells. Cells were stained with 50 μl/well of above antibody cocktail, incubated for 20 min in the dark at room temperature and then washed with 100 μl/well of PBS + 3% FBS. The type (lineage composition) and the size of the colonies formed were assessed by high-throughput flow cytometry (BD FACS Symphony). Colony output was determined using the gating strategy shown in Extended Data Figure 6d. A single cell was defined as giving rise to a colony if the sum of cells detected in the CD45^+^ and GlyA^+^ gates was ≥ 30 cells. Erythroid colonies were identified as CD45^−^GlyA^+^ ≥ 30 cells, Megakaryocyte colonies as CD41^+^ ≥ 30 cells, Myeloid colonies as [(CD45^+^CD14^+^) + (CD45^+^CD15^+^)] ≥ 30 cells, NK colonies as CD45^+^CD56^+^ ≥ 30 cells. All high-throughput screening flow cytometry data was recorded in a blinded way, and correlation between the colony phenotype and originating population was only performed at the final stage. Fisher’s exact test, performed in GraphPad (v7.0), were applied to the proportions of each colony type by stage to determine statistical significance in lineage differentiation potential with development.

### Library Preparation and Sequencing

For the droplet-encapsulation scRNA-seq experiments, 7000 live, single, CD45^+^ or CD45^−^ FACS-isolated cells were loaded on to each lane of the Chromium Controller (10x Genomics). Single cell cDNA synthesis, amplification and sequencing libraries were generated using the Single Cell 3’ Reagent Kit as per the 10x Genomics protocol. The libraries from eight loaded channels from each fetal sample were multiplexed together and sequenced on an Illumina HiSeq 4000. The libraries were distributed over eight lanes per flow cell and sequenced using the following parameters: Read1: 26 cycles, i7: 8 cycles, i5: 0 cycles; Read2: 98 cycles to generate 75bp paired end reads.

For the plate-based scRNA-seq experiments, a slightly modified Smart-seq2 protocol was used as previously described^46^. A total of 45 × 96 well plates (4320 single cells) were analysed from three fetal samples. After cDNA generation, libraries were prepared (384 cells per library) using the Illumina Nextera XT kit. Index v2 sets A, B, C and D were used per library to barcode each cell for multiplexing. Each library was sequenced (384 cells) per lane at a sequencing depth of 1-2 million reads per cell on HiSeq 4000 using v4 SBS chemistry to create 75bp paired end reads.

For the mini bulk RNA-seq experiments, each cell lysate was transferred into a 96-well lo-bind plate then processed using the same modified Smart-seq2 protocol as described above. After cDNA generation, libraries were prepared using the Illumina NexteraXT kit with Index v2 set A to barcode each mini bulk library for multiplexing. All libraries were sequenced on one lane of an Illumina HiSeq 4000 using v4 SBS chemistry to create 75bp paired end reads and aiming to achieve a depth of 10 million reads per library.

### Immunohistochemistry

Formalin fixed, paraffin embedded blocks of fetal livers aged 6PCW, 8PCW, 10PCW and 13PCW were obtained from the HDBR. Each was sectioned at 4μm thickness onto APES-coated slides. Sections were dewaxed for 5 minutes in Xylene (Fisher Chemical) then rehydrated through graded ethanol (99%, 95% and 70%; Fisher Chemical) and washed in running water. Sections were treated with hydrogen peroxide block (1%v/v in water; Sigma) for 10 minutes and rinsed in tap water prior to antigen retrieval. Citrate antigen retrieval was used for all sections. Citrate buffer, pH6 was used with pressure heating for antigen retrieval, and then slides placed in TBS, pH7.6 for 5 minutes prior to staining. Staining was done using the Vector Immpress Kit (Vector Laboratories). Sections were blot dried and blocked sequentially with 2.5% normal horse serum, avidin (Vector Laboratories) and then biotin (Vector Laboratories) for 10 minutes each and blot dried in between. Sections were incubated for 60 minutes with primary antibody diluted in TBS pH7.6 (see Supplementary Table 4 for antibody details). Slides were washed twice in TBS pH7.6 for 5 minutes each before incubation for 30 minutes with the secondary antibody supplied with the kit. Slides were washed twice in TBS pH7.6 for 5 minutes each, and developed using peroxidase chromogen DAB. Sections were counterstained in Mayer’s Haematoxylin for 30 seconds, washed and put in scots tap water for 30 seconds. Slides were dehydrated through graded ethanol (70% to 99%) and then placed in Xylene prior to mounting with DPX (Sigma-Aldrich). Sections were imaged on a Nikon Eclipse 80i microscope using NIS-Elements Fv4.

### Alignment, quantification and quality control of scRNA-seq data

Droplet-based (10x) sequencing data was aligned and quantified using the Cell Ranger Single-Cell Software Suite (version 2.0.2, 10x Genomics Inc) using the GRCh38 human reference genome (official Cell Ranger reference, version 1.2.0). Smart-seq2 sequencing data was aligned with *STAR* (version 2.5.1b), using the STAR index and annotation from the same reference as the 10x data. Gene-specific read counts were calculated using *htseq-count* (version 0.10.0). Cells with fewer than 200 detected genes and for which the total mitochondrial gene expression exceeded 20% were removed. Genes that were expressed in fewer than 3 cells were also removed.

### Doublet detection by support vector machine classification

Doublet exclusion was applied using a support vector machine (SVM) algorithm trained on scRNA-seq data and simulated doublets. Doublet data was obtained by randomly choosing pairs of cells and adding their raw counts by gene. Every pair of cells originated from the same lane in order to avoid introducing doublets that are unlikely to exist, *i.e*. between cells that were loaded in different lanes. The SVM was trained in *Python 3.6.3* using the *SVC* (support vector classifier) from the *svm* module in the sklearn package (version 0.19.1). A grid search was applied during the training of the SVM models to optimise its hyperparameters using the *GridSearchCV* class in *sklearn*.

The doublet SVM classifier was compared with the *scrublet* doublet detector and other custom machine learning approaches which use random forest, k-nearest neighbours and logistic regression. The random forest when trained with grid search parameter optimization was overfitting the data predicting close to 0% doublets. The logistic regression was underfitting suggesting that linear classifiers may not be suited for the problem of doublet detection. The k-nearest neighbours while being the closest to *scrublet* in terms of detection mechanism showed high dependency on manual parameter tuning thus making it a subjective method.

The criteria for assessing a doublet detector were: 1) should detect a very low number of doublets or none in the plate-based scRNA-seq data, 2) the identified doublets should not form a compact separate cluster, and 3) doublets should have higher UMI and gene counts relative to the rest of data. Both *scrublet* and the *svm* doublet detectors satisfied the 3 criteria the difference being however that *scrublet* predicted a much lower percentage of doublets (< 1%) while the *svm* detector predicted rates of 4-6 % which are much closer to previously reported values^78^. In addition, we observed that the svm doublet detector had the capacity to remove more of the outliers, which if present, distort diffusion maps.

### Clustering and downstream analysis of scRNA-seq data

Downstream analysis included data normalisation (*NormalizeData*, LogNormalize method, scaling factor 10000), data feature scaling (*ScaleData*), variable gene detection (*FindVariableGenes*), pca (*RunPCA*, from variable genes) and Louvain graph-based clustering (*FindClusters*, data dimensionality reduction using PCA, clustering resolution manually tuned) performed using the R package Seurat (version 2.3.4). Cluster cell identity was assigned by manual annotation using known marker genes and computed differentially expressed genes (DEGs) using *FindAllMarkers* function in *Seurat* package (Wilcoxon rank sum test, *p*-values adjusted for multiple testing using the Bonferroni correction). For computing DEGs all genes were probed provided they were expressed in at least 25% of cells in either of the two populations compared and the expression difference on a log scale was at least 0.25. Manual annotation was performed iteratively, which included validating proposed cell labels with known markers and further investigating clusters whose gene signatures indicated additional diversity.

Clustering and cell type assignment for fetal liver data was assessed using two additional clustering methods: Agglomerative clustering (with Ward linkage and Euclidean affinity) and Gaussian mixture (*AgglomerativeClustering* class from *cluster* module and *GaussianMixture* from *mixture* module in *sklearn* version 0.19.1 Python 3.6.3). Consensus agreement between the 3 clustering methods was measured by Rand index and adjusted mutual information implemented in the *metrics* module in *sklearn* package. The rand indices and adjusted mutual information values of the Louvain clustering method vs the Agglomerative and Gaussian Mixture clustering method were both 0.84 and 0.83 respectively.

After annotation was completed, a cell type classifier was built by training an SVM on labelled fetal liver scRNA-seq data with grid search for parameter optimization. The SVM was previously compared in terms of accuracy and recall with a random forest and logistic regression classifiers trained on same data. Out of the 3 classifiers the SVM was chosen due to showing a mean accuracy and recall of 95%. Random forest showed 94% for both accuracy and recall. Logistic regression with both l1 and l2 regularization showed 92% accuracy/recall. The lower performance of logistic regression classifiers relative to the other 2 was attributed to lower ability of differentiating some of the early progenitors and cell types within the myeloid compartment. The classifier was used for automatic annotation of the Smart-seq2 data sets to allow identification of biologically meaningful clusters and DEG computation.

Data generated from fetal skin and kidney was pre-processed, normalised, clustered and manually annotated, in parallel with, and using the same pipeline as, the liver data. Skin and kidney data were combined using the *MergeSeurat* function. Clusters characterised by differentially expressed immune gene markers were extracted from the NLT dataset for subsequent comparative analysis with liver-derived immune populations. Human cord blood and adult bone marrow datasets were downloaded from Human Cell Atlas data portal (https://preview.data.humancellatlas.org/). These were processed using the same approach as described above, followed by manual annotation. Alignment of datasets differing in sequencing technology and batch correction were done by canonical correlation analysis (*runCCA*, *Seurat*).

Dimensionality reduction methods included tSNE (Seurat, computed from the first 20 PCs, Barnes-Hut fast computation), UMAP (Python UMAP package, 5 nearest neighbours, correlation metric, minimum distance 0.3, computed from the first 20 PCs), FDG (ForceAtlas2 class from fa2 Python Package, Barnes-Hut implementation for faster computation with theta 0.8, 2000 iterations) and partition-based approximate graph abstraction (PAGA) (*paga* in *scanpy* Python package version 1.2.2). PAGA connectivity scores were used for quantifying cell type similarities across tissues and plotted as heatmap. Development trajectories were inferred by comparing FDG, PAGA and diffusion map plots. Trajectory analysis included computing diffusion map (*scanpy tl.diffmap* with 20 components), pseudotime (*scanpy tl.dpt* setting the earliest known cell type as root) and variable genes across pseudotime.

### Dynamically expressed genes across pseudotime

Genes that vary across pseudotime were calculated using *DifferentialGeneTest* function in Monocle in R (version 2.6.4). This was applied on the entire pseudotime range and also on the pseudotime intervals specific to each cell type in order to avoid limitation to the genes characterised by monotonic changes across the trajectory. Expression of pseudotime variable genes were min-max normalised prior to visualization. The pseudotime variable genes were annotated based on each gene’s involvement in relevant cell-specific functional modules or hallmark functional pathways from MSigDB v6.2, a curated molecular signature database^79^. Transcription factors were marked within the dataset based on the DBD transcription factor prediction database^80^.

Peak expression for each gene over pseudotime was calculated and grouped into ‘Early’, ‘Mid’ or ‘Late’ categories. For visualisation purposes, the resulting gene lists were minimised by ordering them from those present in the most selected functional pathways to least, as well as ensuring coverage across pseudotime. The top 30 genes in each list were displayed using the *DoHeatmap* function of the Seurat package and the full pseudotime gene list is available in the interactive files accompanying diffusion maps.

### Visualisation by animated force-directed graph representation

The FDG animation was created using an in-house modified version of the ForceAtlas2 class in *fa2* Python package by saving all the intermediate states (published version only outputs the final state and discards all intermediates). The FDG coordinates at each iteration were plotted and the resulting graphs were assembled in a mp4 video format using *VideoWriter* in *cv2* (version 3.3.1) Python package.

### Differential marker gene extraction and validation

Differential marker gene validation was done using a random forest classifier (*RandomForestClassifier* class in *ensemble* module of *sklearn* Python package, with 100 estimators, *gini* split criterion, maximum features 15 and maximum depth of 10 splits). To determine whether tissue-related transcriptome variations were present in equivalent immune populations between liver, skin and kidney, each equivalent population was taken in turn and grouped according to its tissue of origin. Seurat *FindMarkers* function was then applied in a pair-wise manner between each tissue subset to produce a cell type-specific list of genes marking each tissue subset. These were investigated in turn for biological relevance, with representative markers displayed using *VlnPlot* function of Seurat.

### Primary immunodeficiency gene list curation

Disease and genetic deficiency information was extracted from Picard et al^15^ and manually annotated to include HGNC symbol names for each disease-associated genetic defect for subsequent correlation with the liver dataset. Diseases implicated in PID were divided according to the International Union of Immunological Societies (IUIS) major categories and screened across the liver scRNA-seq dataset. 315 unique genes were identified in the dataset from the 354 inborn errors of immunity highlighted in the article. For each disease category a dot plot was generated using Seurat *DotPlot* function and ordered by highest expression across each gene and across each cell type, highlighting those cell types in each disease category that express the highest number of genes associated with a genetic defect.

All graphs presented in the manuscript were plotted using ggplot R package or matplotlib Python package. Statistical significance of changes in cellular composition during development were investigated by applying the Kruskal-Wallis test followed by Dunn’s multiple comparison, using GraphPad Prism (v7.0), to population frequencies across all stages.

### CellPhoneDB Analysis

For the receptor-ligand analysis in Figure 3d, a list of curated receptor-ligand interactions was retrieved from CellPhoneDB (www.CellPhoneDB.org). The interactions whose receptor and ligand genes had nonzero expression in at least 30% of cells within two cell types, respectively, were considered valid. Receptor-ligand interactions between VCAM1^+^ Erythroblastic Island macrophages and the two erythroid (early, mid) populations were listed.

### Whole genome sequencing and fetal cell identification

To identify maternal cells present in our data we combined the information from fetal whole genome DNA sequencing with the single cell RNA-seq data. For each sample we measured the allele frequency in the fetal DNA of SNPs from the 1000 genomes project^81^ falling within exons with a population allele frequency in excess of 1%. We then consider only those SNPs which are homozygous in the fetal DNA for follow up in the scRNA-seq data. A SNP was considered to be homozygous if its allele frequency in the fetal DNA was less than 0.2 or greater than 0.8 and had an FDR adjusted p-value of less than 0.01 under a binomial test for the null hypothesis that the allele frequency in the DNA was in the range [0.3,0.7].

The allele frequency of each of these SNPs with population allele frequency > 1% that are known to be homozygous in the fetal DNA was then measured in each cell in the scRNA-seq data. Any deviations from homozygosity in the RNA-seq data must be a consequence of either sequencing errors, RNA editing, or the genotype of the cell differing from the fetal DNA. For each cell, we calculated the total fraction of reads at the SNPs (selected as described above) that differ from the fetal genotype. We then assume that the genome-wide rate of deviations due to sequencing errors and RNA editing is less than or equal to 2%. For maternal cells, the expected genome wide rate of deviation at these SNPs is equal to half the mean of the population allele frequency at the interrogated SNPs. Finally, for each cell we calculated the posterior probability of the cell being fetal or maternal assuming a binomial distribution with rate 2% for a fetal cell and half the mean of the population allele frequency for the maternal cell and assign a cell as: maternal/fetal if either posterior probability exceeds 99%, ambiguous otherwise. We validated this method using samples for which both the fetal and maternal DNA were available.

### ‘Hyperion’ Imaging mass cytometry (IMC)

Antibodies were conjugated to metals using the Fluidigm MaxPar conjugation kits and the associated method with following modifications; the lanthanides were used at 1.5 mM and washed for a shorter duration (4x 5 minutes) in W buffer in prior to elution. Ultrapure MilliQ water was used throughout for any dilutions and washes. 4 µm Formalin-fixed paraffin-embedded sections obtained from 8 and 15 PCW fetal liver tissue blocks were incubated at 60 °C for 1 hour then dewaxed in Xylene (Fisher). After rehydration through graded alcohols (Fisher) and a 5 minute wash in water, the sections were subjected to Heat-Induced Epitope Retrieval with Citrate buffer (pH- 6.0). Sections were then washed in water and PBS (Gibco) and blocked with 3% BSA (Sigma-Aldrich) for 45 minutes. A mixture of 8 metal-conjugated antibodies diluted in 0.5% BSA (see Supplementary Table 5 for antibody details, was added to the sections for overnight incubation at 4 °C in a humidified chamber. Slides were washed twice in 0.2% Triton X-100 diluted in PBS for 8 minutes and then twice in PBS for 8 minutes.

To counterstain nucleated cells, sections were incubated with 312.5nm (193 Ir) Intercalator-Ir (Fluidigm) for 30 minutes at room temperature. Slides were then washed in water for 5 minutes, and allowed to air-dry at room temperature prior to imaging on the Hyperion imaging mass cytometer. Using expected target cell frequencies from previous fluorescence flow cytometry data, Region of Interest (ROI) size was set to 2.8mm by 3.8mm. The ablation energy was set at 2 db with a laser frequency of 200Hz. Each session of ablation generated a .MCD image file containing information for every panorama and ROI measured whereby each 1µm piece of tissue liberated by the laser was analysed for ionic content on a per channel basis by Time of Flight. Single cell segmentation and feature extraction was performed using CellProfiler (v3.1.5). Nuclei were identified using the “IdentifyPrimaryObjects” module where the input images were the sum of the DNA stained Iridium channels (191 and 193) constructed by the “ImageAfterMath” module. The diameter range set for Nuclei identification was 4-15 pixel units. The “ExpandOrShrink” module was used to grow the nuclear segmentation area by 3 pixels to define the cellular area and the “MeasureObjectIntensity” module was used to determine the mean intensity for each cell object identified.

## Author contributions

M.H. and S.A.T conceived the study. M.H.; S.A.T.; E.L.; R.B. and E.S. designed the experiments. Samples were isolated by R.B. and libraries were prepared by E.S. and L.M. with contributions from D.M.P.; R.V-T; J.P.; J.F. Flow cytometry and FACS experiments were performed by R.B.; E.C.; L.J. and D.M. Imaging mass cytometry experiments were performed by M.A.; B.M.; B.I.; D.M.; and A.F. Cytospins were performed by D.D.; J.F.; and *in vitro* culture differentiation experiments were performed by L.J.; D.M. and E.C. Imaging was performed by B.I.; M.A.; F.G.; Y.G. and A.C., and C.M. and M.A. interpreted immunohistochemistry and developmental pathology sections. M.S.K.; B.L.; O.A.; M.T.; D.D.; T.L.T.; M.S.; O.R-R. and A.R. generated adult and cord blood scRNA-seq datasets. D.M.P. performed the computational analysis with contributions from K.G.; M.E.; K.P.; M.Y.and J.B. M.H. interpreted the data with contributions from D.M.P.; R.B.; B.G.; E.L.; I.R.; A.R.; E.C; A-C.V; R.R.; E.P.; M.M.; J.P.; A.F. and P.V. M.H. wrote the manuscript, with contributions from L.J.; R.B.; E.S.; D.M.P; B.G.; E.L.; I.R; A-C. V. and A.R. M.H.; S.A.T.; S.B.; and E.L. co-directed the study. All authors read and accepted the manuscript.

## Acknowledgements

We thank the Newcastle University Flow Cytometry Core Facility, Bioimaging Core Facility, Genomics Facility, NUIT for technical assistance, School of Computing for access to the High-Performance Computing Cluster, Newcastle Molecular Pathology Node Proximity Lab, Alison Farnworth for clinical liaison, Sophie Hambleton for primary immunodeficiency expertise, Helen Chen for immunohistochemistry assistance. The human embryonic and fetal material was provided by the Joint MRC / Wellcome (MR/R006237/1) Human Developmental Biology Resource (www.hdbr.org).

## Data availability

Our fetal liver dataset can be explored using an interactive web portal through the following weblink: http://developmentcellatlas.ncl.ac.uk/datasets/liver_10x

## Code availability

All scripts are available on https://github.com/haniffalab

---

**Extended Data Figure 1: Single cell transcriptome map of fetal liver. a**, UMAP visualisation (left) and violin plots (right) of n=13 displaying fetal livers grouped by stage and coloured by gender; *XIST* (green) and *RSP4Y1* (purple) expression marks female and male samples respectively. **b**, UMAP visualisation of additional clustering algorithms to assess agreement between methods evaluated by Rand index and Adjusted Mutual Information values. **c**, UMAP visualisation of fetal liver cells coloured by the assigned annotation, and by developmental stage (6-9 PCW, 9-12 PCW, 12-15 PCW, and 15-18 PCW). **d,** UMAP visualisation of 1, 206 fetal liver cells (n=2) profiled using Smart-seq2. **e,** Frequency (mean +/− SEM) of B cells in whole liver detected by scRNA-seq from **1c** (left) and in the CD34^−^ cells detected by flow cytometry (right) from 6-19 PCW (n=53). **f,** Ratio of Mega-erythroid cells (Megakaryocytes and erythroid cells from Figure 1d) cells to Immune (from Figure 1d) (mean +/− SEM). * p < 0.05; *** p < 0.005; **** p < 0.001

**Extended Data Figure 3: Fetal liver and NLT haematopoiesis. a**, AGA analysis of fetal liver HSC/MPP, erythroid, megakaryocyte and mast cell lineages from **Figure 3a**. **b**, Violin plots showing increasing haemoglobin gene expression values of erythroid cell stages in liver (n=13) and skin (n=8). **c,** Dynamically expressed pseudotime genes during erythroid, megakaryocyte and mast cell development, transcription factors marked with *. **d,** AGA connectivity scores of HSC/MPP, erythroid, megakaryocyte and mast cell lineages between fetal liver, skin and kidney from samples F1 to F10. **e,** FDG visualisation of F1-F10 fetal liver, skin and kidney HSC/MPP, MEMP, erythroid, megakaryocyte and mast cell lineages. **f,** Cell cycle analysis (G1, G2M and S phase; bar plots) of HSC/MPP and lymphoid cell types from F1-F10 liver, skin and kidney samples and the % cells positive for *MKI67* expression (% above plots) (red = liver, blue = skin, green = kidney).

**Extended Data Figure 4: Lymphoid lineages in fetal liver and NLT. a**, AGA analysis of fetal liver HSC/MPP and lymphoid subsets from **1c** over three stages (6-9 PCW, 12-15 PCW and 15-18 PCW). Violin plots (**b**) and feature plots (**c**) showing expression values of V(D)J and select genes over gestation for early lymphoid/T lymphocyte cluster (n=13). **d,** AGA connectivity scores of HSC/MPP and lymphoid cell types from F1-F10 fetal liver, skin and kidney. **e,** Violin plots showing expression values of mono-NK and mac-NK cells in F1-F10 fetal liver and skin. **f,** Violin plots showing expression values of genes expressed in ILC precursors from F1-F10 fetal liver, skin and kidney.

**Extended Data Figure 5: Tissue signatures in developing myeloid cells. a**, 3D diffusion map of fetal liver HSC/MPP and myeloid cells from **1c**. **b,** AGA connectivity scores of HSC/MPP and myeloid cells from F1-F10 fetal liver, skin and kidney. **c,** Violin plots showing expression values of select genes in pDC precursors from F1-F10 fetal liver and skin.

**Extended Data Figure 6: HSC/MPP differentiation potential by gestation. a**, 3D diffusion map, **b**, AGA and **c**, FDG visualisation of fetal liver HSC/MPP and progenitor cells from **1c**. **d,** FDG projection of 349 scRNA-seq profiled cells from FACS gates in **6c** with 10x dataset of HSC/MPP and early progenitors from **1c**. **e,** Bar plot showing the total % of colonies (undifferentiated), one (unilineage), two (bilineage) or more (multilineage) from ‘HSC pool’ gate by gestation stage (n=5). **f,** Bar plots showing total % of NK and erythroid colonies from ‘HSC pool’ gate by gestation stage (n=5) **g,** Bar plot showing total % colonies cultured from ‘HSC pool’ gate without MS5 stroma layer. **h**, Percentage of HSC/MPP and early progenitors in fetal liver (n=13), cord blood (n=8) and adult bone marrow (n=8) expressing MKI67. ** p < 0.01; *** p < 0.005

**Supplementary Data Figure 1: FACS gating strategy for scRNA-seq analysis. a**, FACS gating strategy used to isolate sort cells for droplet-(10x) and plate-based scRNA-seq for F1-F15. **b**, FACS gating strategy used to isolate cells for cytospins and mini bulk RNA-seq (n=3). **c,** Flow cytometry gating strategy used to identify the colonies that differentiated from single cell *in vitro* differentiation culture in **Figure 6c**.

**Supplementary Data Figure 2: Expression of known PID genes in fetal liver.** Dot plot showing relative expression of genes known to be associated with PID in fetal liver cell types from Figure 1c. Dot plots according to major PID disease categories. Gene expression frequency indicated by spot size and level by colour intensity.

**Supplementary Data Figure 3: Transcriptome validation of cell types and fetal skin and kidney cells. a,** Dot plot characterising marker gene expression 18 fetal liver cell types. Gene expression frequency indicated by spot size and level by colour intensity **b,** UMAP visualisation of haematopoietic cells from F1-F10 fetal skin and kidney.

## References

1. Jagannathan-Bogdan, M. & Zon, L. I. Hematopoiesis. Development 140, 2463 (2013).

2. Parekh, C. & Crooks, G. M. Critical Differences in Hematopoiesis and Lymphoid Development Between Humans and Mice. J. Clin. Immunol. 33, 711–715 (2013).

3. Ivanovs, A. et al. Human haematopoietic stem cell development: from the embryo to the dish. Development 144, 2323–2337 (2017).

4. Christensen, J. L., Wright, D. E., Wagers, A. J. & Weissman, I. L. Circulation and Chemotaxis of Fetal Hematopoietic Stem Cells. PLOS Biol. 2, e75 (2004).

5. Mikkola, H. K. A. & Orkin, S. H. The journey of developing hematopoietic stem cells. Development 133, 3733 (2006).

6. Wiemels, J. Perspectives on the Causes of Childhood Leukemia. Chem. Biol. Interact. 196, 10.1016/j.cbi.2012.01.007 (2012).

7. Roberts, I. & Chakravorty, S. Neonatal haematology. Postgrad. Haematol. 870–884 (2015).

8. Holt, P. G. & Jones, C. A. The development of the immune system during pregnancy and early life. Allergy 55, 688–697 (2001).

9. Tavian, M. & Peault, B. Embryonic development of the human hematopoietic system. Int. J. Dev. Biol. 49, 243–250 (2003).

10. Julien, E., El Omar, R. & Tavian, M. Origin of the hematopoietic system in the human embryo. FEBS Lett. 590, 3987–4001 (2016).

11. Kashem, S. W., Haniffa, M. & Kaplan, D. H. Antigen-Presenting Cells in the Skin. Annu. Rev. Immunol. 35, 469–499 (2017).

12. Mass, E. et al. Specification of tissue-resident macrophages during organogenesis. Science (2016). doi:10.1126/science.aaf4238

13. Butler, A., Hoffman, P., Smibert, P., Papalexi, E. & Satija, R. Integrating single-cell transcriptomic data across different conditions, technologies, and species. Nat. Biotechnol. 36, 411 (2018).

14. Ohls, R. K. et al. Neutrophil Pool Sizes and Granulocyte Colony-Stimulating Factor Production in Human Mid-Trimester Fetuses. Pediatr. Res. 37, 806 (1995).

15. Picard, C. et al. International Union of Immunological Societies: 2017 Primary Immunodeficiency Diseases Committee Report on Inborn Errors of Immunity. J. Clin. Immunol. 38, 96–128 (2018).

16. Bhogal, R. H. et al. Isolation of Primary Human Hepatocytes from Normal and Diseased Liver Tissue: A One Hundred Liver Experience. PLoS ONE 6, e18222 (2011).

17. Green, C. J. et al. The isolation of primary hepatocytes from human tissue: optimising the use of small non-encapsulated liver resection surplus. Cell Tissue Bank. 18, 597–604 (2017).

18. MacParland, S. A. et al. Single cell RNA sequencing of human liver reveals distinct intrahepatic macrophage populations. Nat. Commun. 9, 4383 (2018).

19. Tusi, B. K. et al. Population snapshots predict early haematopoietic and erythroid hierarchies. Nature 555, 54 (2018).

20. Zheng, S., Papalexi, E., Butler, A., Stephenson, W. & Satija, R. Molecular transitions in early progenitors during human cord blood hematopoiesis. Mol. Syst. Biol. 14, e8041 (2018).

21. Gentek, R. et al. Hemogenic Endothelial Fate Mapping Reveals Dual Developmental Origin of Mast Cells. Immunity 48, 1160–1171.e5 (2018).

22. Li, Z. et al. Adult Connective Tissue-Resident Mast Cells Originate from Late Erythro-Myeloid Progenitors. Immunity 49, 640–653 (2018).

23. Klei, T. R. L., Meinderts, S. M., van den Berg, T. K. & van Bruggen, R. From the Cradle to the Grave: The Role of Macrophages in Erythropoiesis and Erythrophagocytosis. Front. Immunol. 8, 73 (2017).

24. Chow, A. et al. CD169+ macrophages provide a niche promoting erythropoiesis under homeostasis and stress. Nat. Med. 19, 429 (2013).

25. Kessel, K. U. et al. Emergence of CD43-Expressing Hematopoietic Progenitors from Human Induced Pluripotent Stem Cells. Transfus. Med. Hemotherapy 44, 143–150 (2017).

26. Seu, K. G. et al. Unraveling Macrophage Heterogeneity in Erythroblastic Islands. Front. Immunol. 8, 1140 (2017).

27. Sadahira, Y., Yoshino, T. & Monobe, Y. Very late activation antigen 4-vascular cell adhesion molecule 1 interaction is involved in the formation of erythroblastic islands. J. Exp. Med. 181, 411 (1995).

28. Ihanus, E., Uotila, L. M., Toivanen, A., Varis, M. & Gahmberg, C. G. Red-cell ICAM-4 is a ligand for the monocyte/macrophage integrin CD11c/CD18: characterization of the binding sites on ICAM-4. Blood 109, 802 (2007).

29. An, X. et al. Global transcriptome analyses of human and murine terminal erythroid differentiation. Blood 123, 3466 (2014).

30. Gautier, E.-F. et al. Comprehensive Proteomic Analysis of Human Erythropoiesis. Cell Rep. 16, 1470–1484 (2016).

31. Okada, Y. et al. Homeodomain proteins MEIS1 and PBXs regulate the lineage-specific transcription of the platelet factor 4 gene. Blood 101, 4748 (2003).

32. Dedhia, P., Kambayashi, T. & Pear, W. S. Notch2 paves the way to mast cells by Hes1 and Gata3. Proc. Natl. Acad. Sci. 105, 7629 (2008).

33. Cheadle, C. et al. Erythroid-Specific Transcriptional Changes in PBMCs from Pulmonary Hypertension Patients. PLOS ONE 7, e34951 (2012).

34. Ebert, B. L. et al. An Erythroid Differentiation Signature Predicts Response to Lenalidomide in Myelodysplastic Syndrome. PLOS Med. 5, e35 (2008).

35. Gale, R. P. Development of the immune system in human fetal liver. in Fetal liver transplantation (eds. Touraine, J.-L., Gale, R. P. & Kochupillai, V.) 45–56 (Springer Netherlands, 1987). doi:10.1007/978-94-009-3365-1_6

36. Phillips, J. H. et al. Ontogeny of human natural killer (NK) cells: fetal NK cells mediate cytolytic function and express cytoplasmic CD3 epsilon, delta proteins. J. Exp. Med. 175, 1055 (1992).

37. Forkel, M. et al. Composition and functionality of the intrahepatic innate lymphoid cell-compartment in human nonfibrotic and fibrotic livers. Eur. J. Immunol. 47, 1280–1294 (2017).

38. Barcena, A. et al. Phenotypic and functional analysis of T-cell precursors in the human fetal liver and thymus: CD7 expression in the early stages of T- and myeloid-cell development. Blood 82, 3401 (1993).

39. Haynes, B. F. & Heinly, C. S. Early human T cell development: analysis of the human thymus at the time of initial entry of hematopoietic stem cells into the fetal thymic microenvironment. J. Exp. Med. 181, 1445–1458 (1995).

40. Sánchez, M. J., Spits, H., Lanier, L. L. & Phillips, J. H. Human natural killer cell committed thymocytes and their relation to the T cell lineage. J. Exp. Med. 178, 1857 (1993).

41. Darrasse-Jèze, G., Marodon, G., Salomon, B. L., Catala, M. & Klatzmann, D. Ontogeny of CD4^+^CD25^+^ regulatory/suppressor T cells in human fetuses. Blood 105, 4715 (2005).

42. Wucherpfennig, K. W. et al. Structural requirements for binding of an immunodominant myelin basic protein peptide to DR2 isotypes and for its recognition by human T cell clones. J. Exp. Med. 179, 279 (1994).

43. Li, N. et al. Mass cytometry reveals innate lymphoid cell differentiation pathways in the human fetal intestine. J. Exp. Med. (2018). doi:10.1084/jem.20171934

44. Spits, H. et al. Innate lymphoid cells — a proposal for uniform nomenclature. Nat. Rev. Immunol. 13, 145 (2013).

45. Chen, L. et al. CD56 Expression Marks Human Group 2 Innate Lymphoid Cell Divergence from a Shared NK Cell and Group 3 Innate Lymphoid Cell Developmental Pathway. Immunity 49, 464–476.e4 (2018).

46. Villani, A.-C. et al. Single-cell RNA-seq reveals new types of human blood dendritic cells, monocytes, and progenitors. Science 356, (2017).

47. Itoh, K. & Hirohata, S. The role of IL-10 in human B cell activation, proliferation, and differentiation. J. Immunol. 154, 4341 (1995).

48. Stremmel, C. et al. Yolk sac macrophage progenitors traffic to the embryo during defined stages of development. Nat. Commun. 9, 75 (2018).

49. Ginhoux, F. & Jung, S. Monocytes and macrophages: developmental pathways and tissue homeostasis. Nat. Rev. Immunol. 14, 392 (2014).

50. Merad, M., Sathe, P., Helft, J., Miller, J. & Mortha, A. The Dendritic Cell Lineage: Ontogeny and Function of Dendritic Cells and Their Subsets in the Steady State and the Inflamed Setting. Annu. Rev. Immunol. 31, 563–604 (2013).

51. Murphy, T. L. et al. Transcriptional Control of Dendritic Cell Development. Annu. Rev. Immunol. 34, 93–119 (2016).

52. Sathe, P., Vremec, D., Wu, L., Corcoran, L. & Shortman, K. Convergent differentiation: myeloid and lymphoid pathways to murine plasmacytoid dendritic cells. Blood 121, 11 (2013).

53. Helft, J. et al. Dendritic Cell Lineage Potential in Human Early Hematopoietic Progenitors. Cell Rep. 20, 529–537 (2017).

54. Rodrigues, P. F. et al. Distinct progenitor lineages contribute to the heterogeneity of plasmacytoid dendritic cells. Nat. Immunol. 19, 711–722 (2018).

55. Yona, S. et al. Fate Mapping Reveals Origins and Dynamics of Monocytes and Tissue Macrophages under Homeostasis. Immunity 38, 79–91 (2013).

56. T’Jonck, W., Guilliams, M. & Bonnardel, J. Niche signals and transcription factors involved in tissue-resident macrophage development. Spec. Issue Tissue Macrophage Compend. 330, 43–53 (2018).

57. van de Laar, L. et al. Yolk Sac Macrophages, Fetal Liver, and Adult Monocytes Can Colonize an Empty Niche and Develop into Functional Tissue-Resident Macrophages. Immunity 44, 755–768 (2016).

58. Guilliams, M. et al. Alveolar macrophages develop from fetal monocytes that differentiate into long-lived cells in the first week of life via GM-CSF. J. Exp. Med. 210, 1977 (2013).

59. McGovern, N. et al. Human Dermal CD14+ Cells Are a Transient Population of Monocyte-Derived Macrophages. Immunity 41, 465–477 (2014).

60. McGovern, N. et al. Human fetal dendritic cells promote prenatal T-cell immune suppression through arginase-2. Nature 546, 662 (2017).

61. Grün, D. et al. De Novo Prediction of Stem Cell Identity using Single-Cell Transcriptome Data. Cell Stem Cell 19, 266–277 (2016).

62. Velten, L. et al. Human haematopoietic stem cell lineage commitment is a continuous process. Nat. Cell Biol. 19, 271 (2017).

63. Belluschi, S. et al. Myelo-lymphoid lineage restriction occurs in the human haematopoietic stem cell compartment before lymphoid-primed multipotent progenitors. Nat. Commun. 9, 4100 (2018).

64. Laurenti, E. et al. The transcriptional architecture of early human hematopoiesis identifies multilevel control of lymphoid commitment. Nat. Immunol. 14, 756 (2013).

65. Morrison, S. J., Hemmati, H. D., Wandycz, A. M. & Weissman, I. L. The purification and characterization of fetal liver hematopoietic stem cells. Proc. Natl. Acad. Sci. 92, 10302 (1995).

66. Bowie, M. B. et al. Hematopoietic stem cells proliferate until after birth and show a reversible phase-specific engraftment defect. J. Clin. Invest. 116, 2808–2816 (2006).

67. Copley, M. R. et al. The Lin28b–let-7–Hmga2 axis determines the higher self-renewal potential of fetal haematopoietic stem cells. Nat. Cell Biol. 15, 916 (2013).

68. Daum, L. M., Sklar, L. R. & Mehregan, D. R. Blueberry muffin rash secondary to hereditary spherocytosis. Cutis 101, 111–114 (2018).

69. Schofield, J. K., Shun, J. L. K., Cerio, R. & Grice, K. Cutaneous extramedullary hematopoiesis with a preponderance of atypical megakaryocytes in myelofibrosis. J. Am. Acad. Dermatol. 22, 334–337 (1990).

70. Kingsley, P. D., Malik, J., Fantauzzo, K. A. & Palis, J. Yolk sac–derived primitive erythroblasts enucleate during mammalian embryogenesis. Blood 104, 19 (2004).

71. Lefrançais, E. et al. The lung is a site of platelet biogenesis and a reservoir for haematopoietic progenitors. Nature 544, 105 (2017).

72. Lysakova-Devine, T. & O’Farrelly, C. Tissue-specific NK cell populations and their origin. J. Leukoc. Biol. 96, 981–990 (2014).

73. Huang, Y., Mao, K. & Germain, R. N. Thinking differently about ILCs—Not just tissue resident and not just the same as CD4+ T-cell effectors. Immunol. Rev. 286, 160–171 (2018).

74. Gerrelli, D., Lisgo, S., Copp, A. J. & Lindsay, S. Enabling research with human embryonic and fetal tissue resources. Development 142, 3073 (2015).

75. Bullen, P. & Wilson, D. The Carnegie staging of human embryos: a practical guide. Mol. Genet. Early Hum. Dev. 27–35 (1997).

76. Hern, W. M. Correlation of fetal age and measurements between 10 and 26 weeks of gestation. Obstet Gynecol 63, (1984).

77. Roy, A. et al. High resolution IgH repertoire analysis reveals fetal liver as the likely origin of life-long, innate B lymphopoiesis in humans. Clin. Immunol. 183, 8–16 (2017).

78. Macosko, E. Z. et al. Highly Parallel Genome-wide Expression Profiling of Individual Cells Using Nanoliter Droplets. Cell 161, 1202–1214 (2015).

79. Subramanian, A. et al. Gene set enrichment analysis: A knowledge-based approach for interpreting genome-wide expression profiles. Proc. Natl. Acad. Sci. 102, 15545 (2005).

80. Zhang, H.-M. et al. AnimalTFDB: a comprehensive animal transcription factor database. Nucleic Acids Res. 40, D144–D149 (2012).

81. The 1000 Genomes Project Consortium et al. A global reference for human genetic variation. Nature 526, 68 (2015).

