## Extended Data Figure 1-6; Extended Data Table 1-3 for "Decoding the development of the blood and immune systems during human fetal liver haematopoiesis"

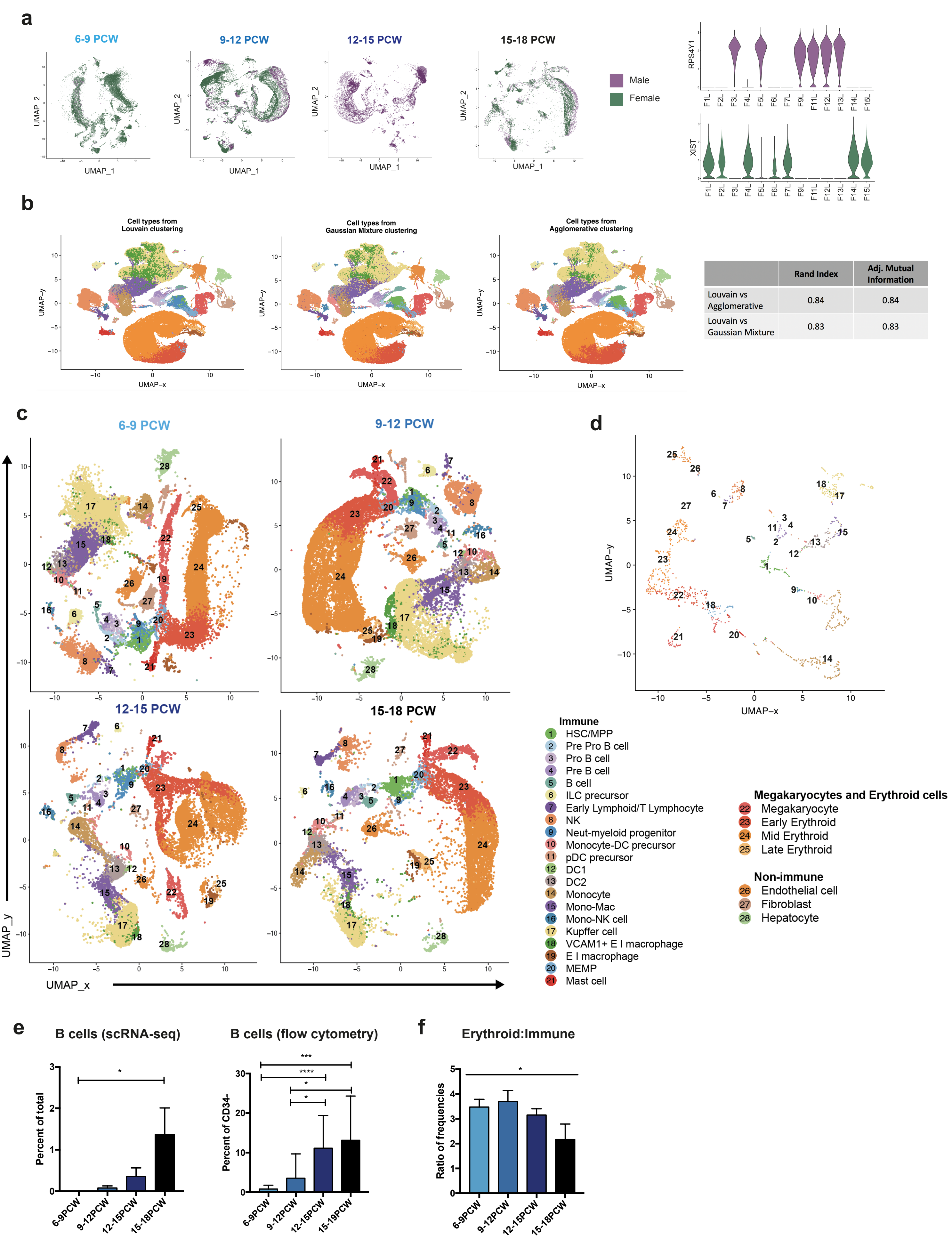


| | Rand Index | Adj. Mutual Information |
|---|---|---|
| Louvain vs Agglomerative | 0.84 | 0.84 |
| Louvain vs Gaussian Mixture | 0.83 | 0.83 |

### Extended Data Figure 3

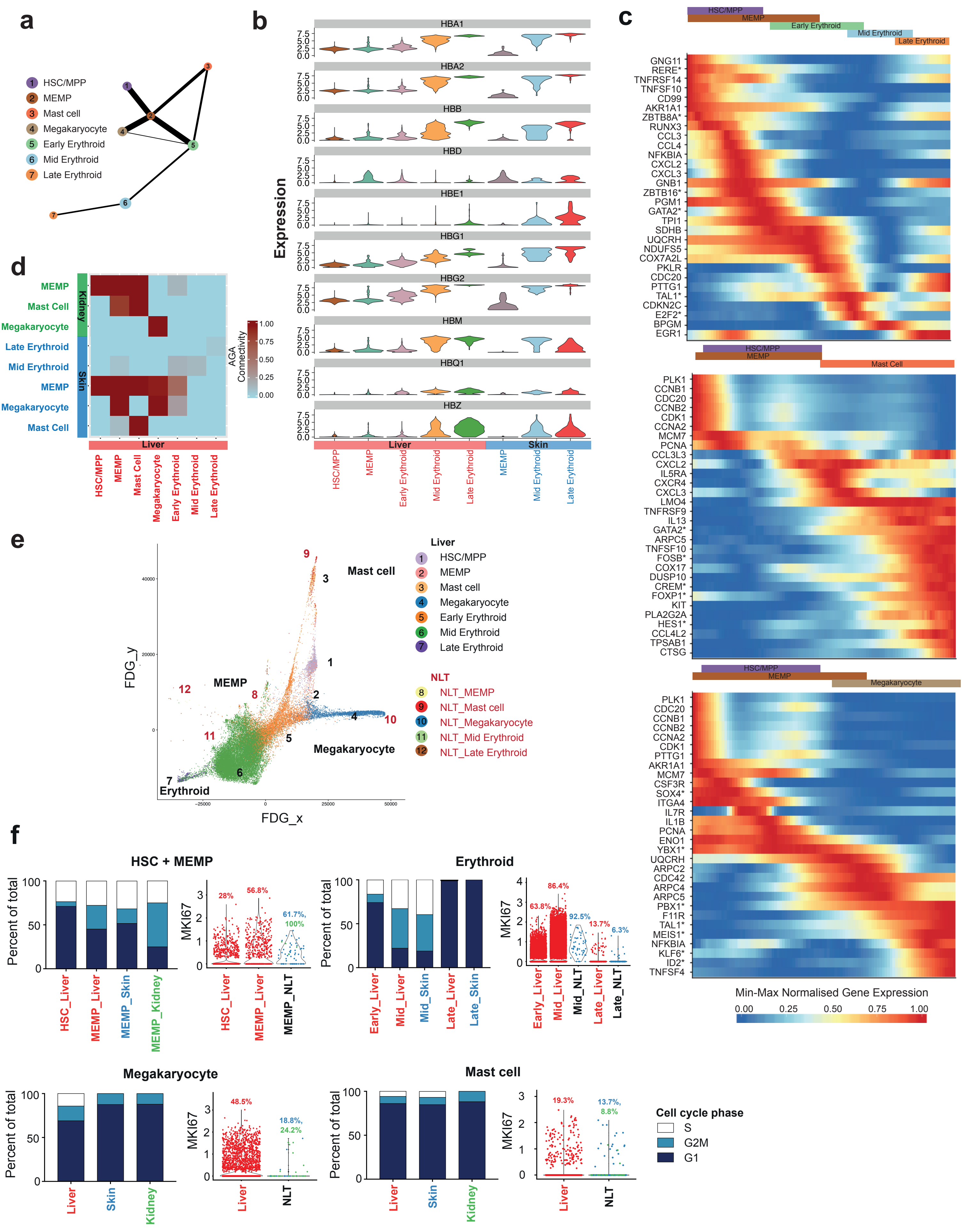

### Extended Data Figure 4

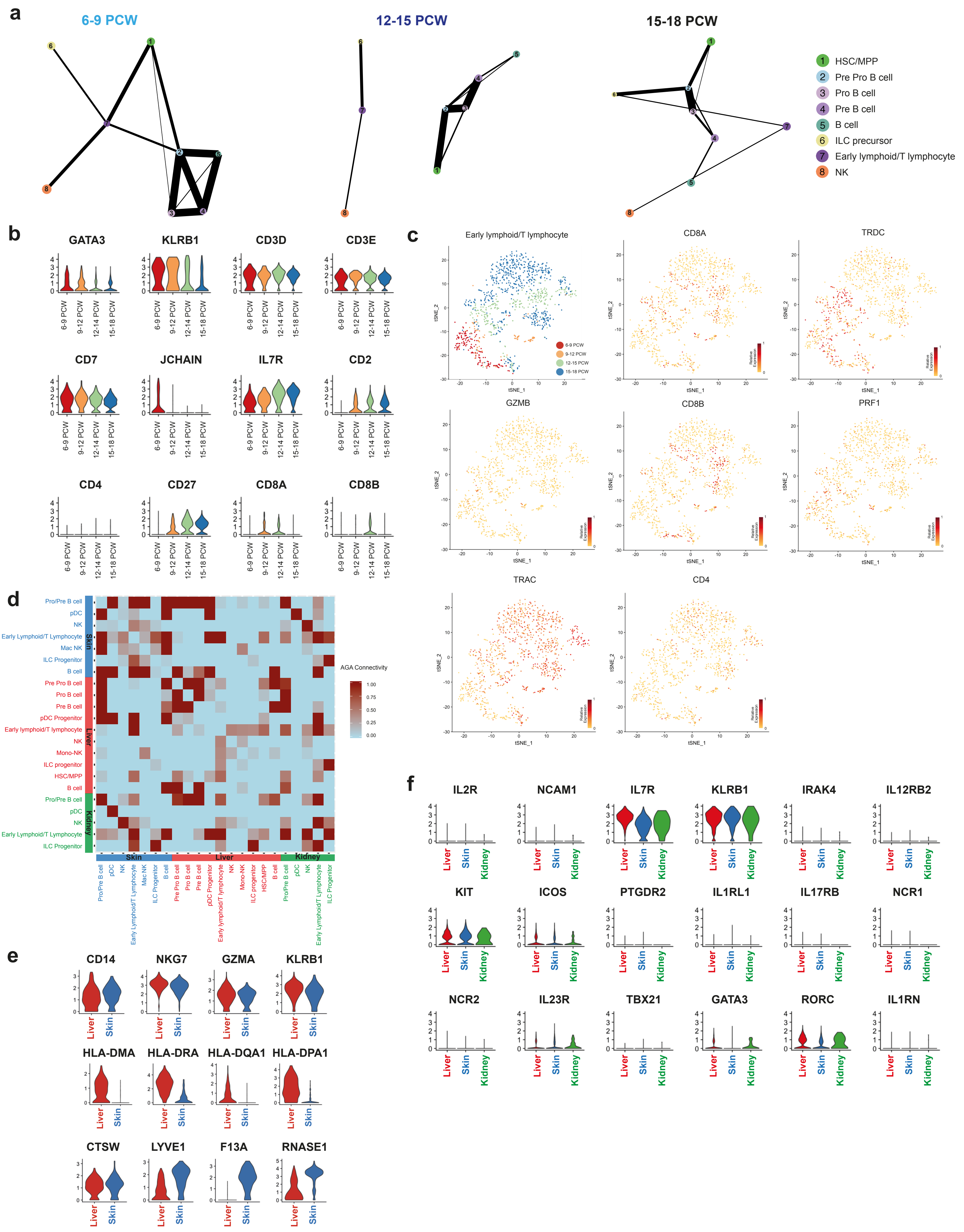

### Extended Data Figure 5

**a**

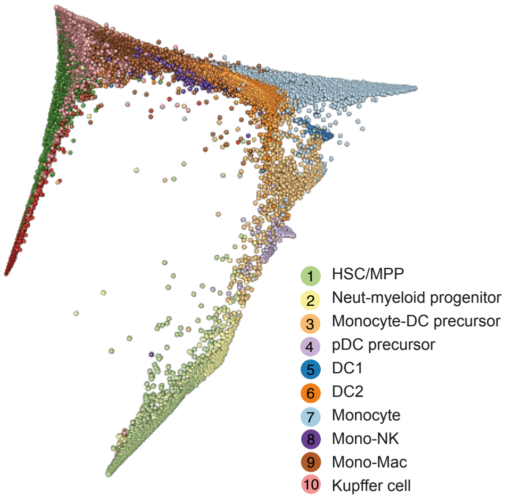


**b**

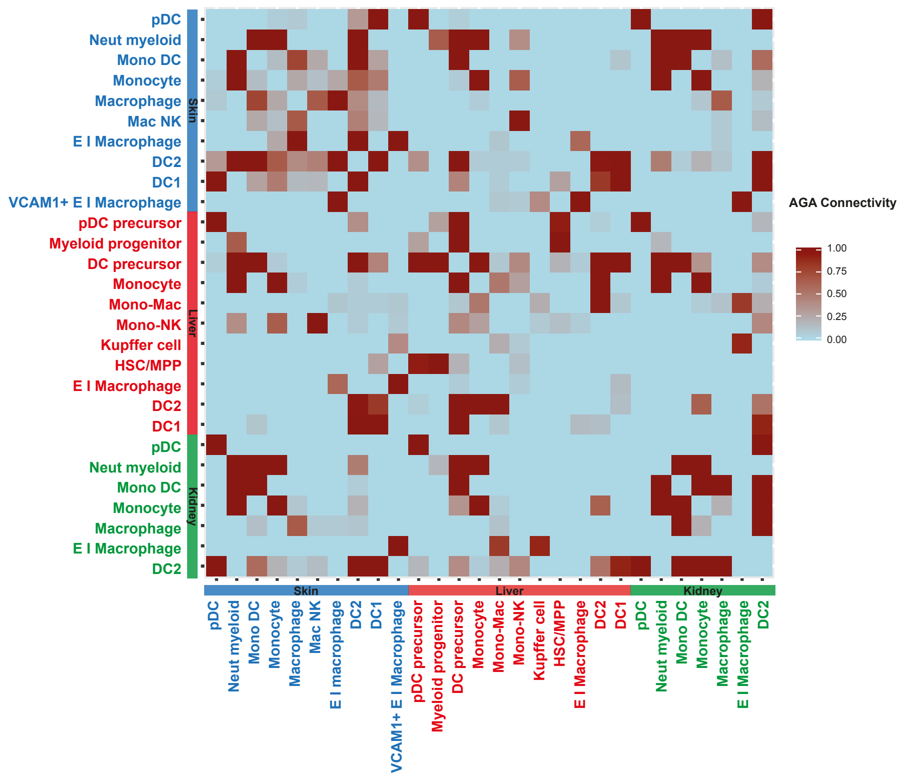


**c**

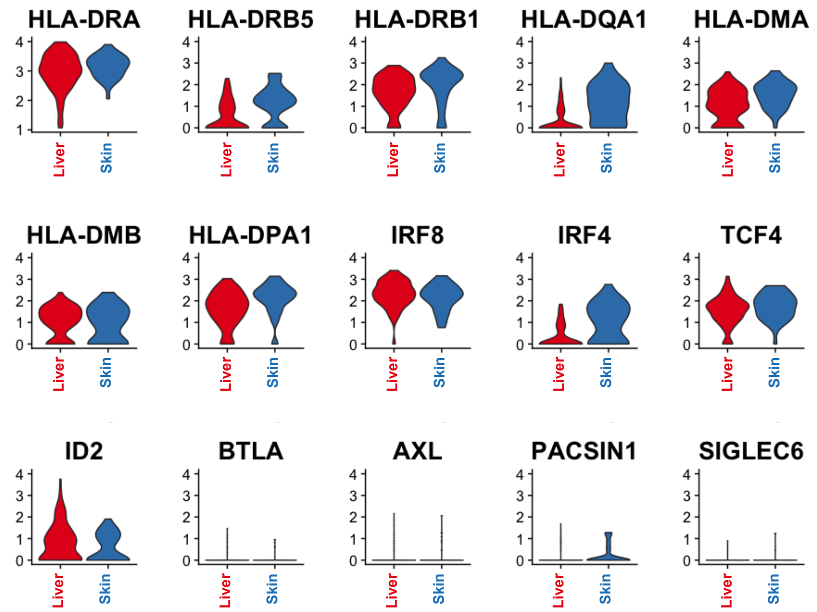

### Extended Data Figure 6

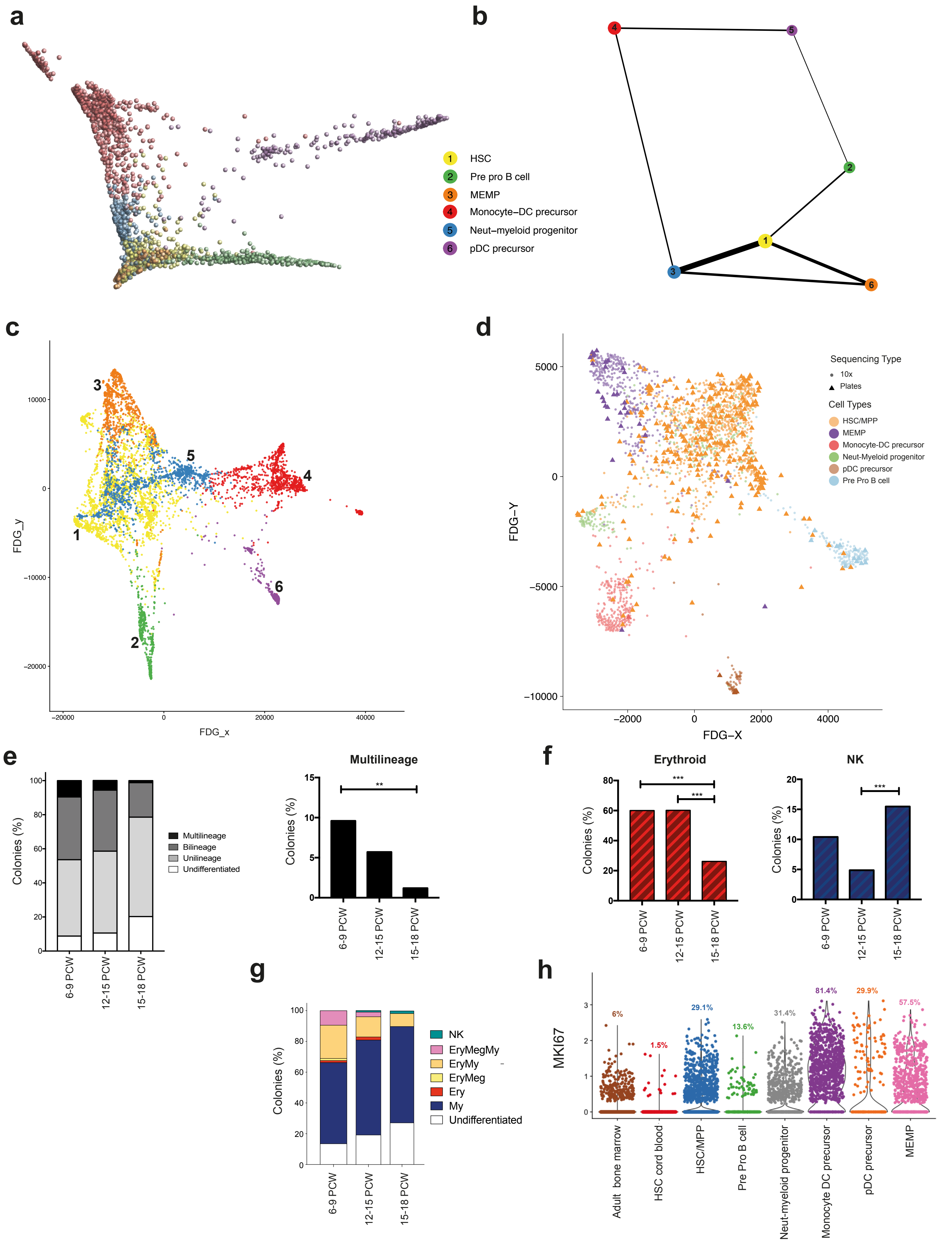

#### Extended Data Table 1. Sample details

| Sample | Type | Age | Stage | Sex | Tissue | Use | scRNA-seq | | |
|---|---|---|---|---|---|---|---|---|---|
| | | | | | | | Gate (sort %) | No. cells | |
| | | | | | | | | 10x* | SS2 |
| F1 | medical | 7 PCW | 6-9 PCW | F | Liver | sR | CD45+ (12.1) | 6,351 | |
| | | | | | | | CD45- (87.9) | 3,620 | |
| | | | | | Skin | sR | CD45+ (3.6) | 2,822 | |
| | | | | | | | CD45- (96.4) | 7,682 | |
| F2 | medical | 7 PCW | 6-9 PCW | F | Liver | sR | CD45+ (5.7) | 6,244 | |
| | | | | | | | CD45- (94.3) | 3,477 | |
| | | | | | Skin | sR | CD45+ (1.4) | 3,492 | |
| | | | | | | | CD45- (98.6) | 4,968 | |
| | | | | | Kidney | sR | CD45+ (1.6) | 1,970 | |
| | | | | | | | CD45- (98.4) | 4,467 | |
| F3 | surgical | 8 PCW | 6-9 PCW | M | Liver | sR | CD45+ (12) | 157 | |
| | | | | | | | CD45- (88) | 1,201 | |
| | | | | | Skin | sR | CD45+ (3.4) | 573 | |
| | | | | | | | CD45- (96.6) | 369 | |
| | | | | | Kidney | sR | CD45+ (6.2) | 345 | |
| | | | | | | | CD45- (93.8) | 736 | |
| F4 | medical | 8 PCW | 6-9 PCW | F | Liver | sR | CD45+ (11.1) | 11,827 | |
| | | | | | | | CD45- (88.9) | 7,317 | |
| | | | | | Skin | sR | CD45+ (3.2) | 2,601 | |
| | | | | | | | CD45- (96.8) | 12,857 | |
| F5 | surgical | 9 PCW | 9-12 PCW | M | Liver | sR | CD45+ (14.5) | 285 | 95 |
| | | | | | | | CD45- (85.5) | 745 | 89 |
| | | | | | | | Live | 1,250 | |
| | | | | | Skin | sR | CD45+ (2.2) | 260 | 93 |
| | | | | | | | CD45- (97.8) | 788 | 93 |
| | | | | | | | Live | 1,103 | |
| | | | | | Kidney | sR | CD45+ (3.2) | 399 | 94 |
| | | | | | | | CD45- (96.8) | 944 | 188 |
| | | | | | | | Live | 1,047 | |
| F6 | medical | 9 PCW | 9-12 PCW | F | Liver | sR | CD45+ (11.1) | 2,713 | |
| | | | | | | | CD45- (88.9) | 3,784 | |
| | | | | | | | Live | 4,923 | |
| F7 | surgical | 9 PCW | 9-12 PCW | F | Liver | sR | CD45+ (12.4) | 12,160 | |
| | | | | | | | CD45- (87.6) | 7,043 | |
| | | | | | Skin | sR | CD45+ (5.2) | 8,712 | |
| | | | | | | | CD45- (94.8) | 6,023 | |
| F8 | surgical | 10 PCW | 9-12 PCW | F | Skin | sR | CD45+ (2.5) | 313 | |
| | | | | | | | CD45- (97.5) | 5,126 | |
| F9 | medical | 11 PCW | 9-12 PCW | M | Liver | sR | CD45+ (11.8) | 3,073 | |
| | | | | | | | CD45- (88.2) | 3,247 | |
| F10 | surgical | 12 PCW | 12-15 PCW | M | Skin | sR | CD45+ (9.3) | 91 | |
| | | | | | | | CD45- (90.7) | 67 | |
| | | | | | Kidney | sR | CD45+ (7.6) | 87 | 92 |
| | | | | | | | CD45- (92.4) | 117 | 138 |
| F11 | medical | 12 PCW | 12-15 PCW | M | Liver | sR | CD45+ (12.1) | 4,719 | |
| | | | | | | | CD45- (87.9) | 5,761 | |
| F12 | medical | 14 PCW | 12-15 PCW | M | Liver | sR | CD45+ (16.9) | 8,678 | |
| | | | | | | | CD45- (83.1) | 4,748 | |
| F13 | medical | 16 PCW | 15-18 PCW | M | Liver | sR | CD45+ (27.2) | 3,039 | |
| | | | | | | | CD45- (72.8) | 4,447 | |
| F14 | medical | 16 PCW | 15-18 PCW | F | Liver | sR, C, M, bR | CD45+ (13.3) | 10,456 | |
| | | | | | | | CD45- (86.7) | 4,197 | |
| F15 | medical | 17 PCW | 15-18 PCW | F | Liver | sR | CD45+ (26.8) | 4,649 | |
| | | | | | | | CD45- (73.2) | 3,658 | |
| F16 | medical | 9 CPW | 6-9 PCW | | Liver | C | | | |
| F17 | surgical | 12 PCW | 12-15 PCW | F | Liver | sR | CD45+ (20) | | 466 |
| | | | | | | | CD45- (80) | | 556 |
| | | | | | Skin | sR | CD45+ (5.6) | | 542 |
| | | | | | | | CD45- (94.4) | | 502 |
| | | | | | Kidney | sR | CD45+ (18.6) | | 240 |
| | | | | | | | CD45- (81.4) | | 373 |
| F18 | medical | 13 PCW | 12-15 PCW | | Liver | C, M | | | |

PCW = post conception weeks; F = female; M = male; sR = single cell RNA-seq; bR = bulk RNA-seq; C = culture; M = morphology by cytospin; I = imaging; * = includes cell numbers from 10x Genomics 3’ and 5’ kits

**Extended Data Table 2. Cell numbers before and after QC**

| PCW | Sample | Liver | | | | Skin | | Kidney | |
|---|---|---|---|---|---|---|---|---|---|
| | | Cell Ranger Cells | Passed QC Cells | Cell Ranger Cells | Passed QC cells | Cell Ranger Cells | Passed QC Cells | Cell Ranger Cells | Passed QC Cells |
| 6-9 | F1 | 7500 | 7486 | 2587* | 2485* | 10505 | 10504 | | |
| | F2 | 9745 | 9721 | | | 8460 | 8460 | 6437 | 6437 |
| | F3 | 1377 | 1358 | | | 970 | 969 | 1084 | 1081 |
| | F4 | 17477 | 17370 | 1867* | 1774* | 15460 | 15458 | | |
| 9-12 | F5 | 1040 | 1030 | | | 1048 | 1048 | 1397 | 1393 |
| | F6 | 6516 | 6497 | | | | | | |
| | F7 | 19323 | 19203 | | | 14740 | 14735 | | |
| | F8 | | | | | 5458 | 5439 | | |
| | F9 | 6333 | 6325 | | | | | | |
| | F10 | | | | | 159 | 158 | 253 | 204 |
| 12-15 | F11 | 15635 | 10480 | 2661* | 2601* | | | | |
| | F12 | 10840 | 10825 | 6136* | 6605* | | | | |
| 15-18 | F13 | 7498 | 7486 | | | | | | |
| | F14 | 13232 | 7900 | 6824* | 6753* | | | | |
| | F15 | 6226 | 6205 | 2223* | 2192* | | | | |

*10x Genomics 5’ kit used

**Extended Data Table 3. Marker gene table**

| Cluster Annotation | Marker Genes |
|---|---|
| Kupffer Cell | VCAM1, CD14, FCGR3A, HMOX1, TIMD4, FOLR2, LGMN, SLC40A1, CETP, MARCO, CD68, FABP3, LIPA, C1QC, C1QB, C1QA, CFP |
| Mono-Mac | RNASE6, CLEC7A, CTSH, TIMD4, FOLR2, SLC40A1, CSTA, CD14, FCGR3A, MS4A7, CD68, MARCO |
| Erythroblastic Island Macrophage | GYPA, CD14, LIPA, FOLR2, FABP3, CD68, MARCO |
| VCAM1+ Erythroblastic Island Macrophage | VCAM1, GYPA, CD14, LIPA, FOLR2, FABP3, CD68, MARCO |
| DC2 | CLEC10A, RNASE6, MNDA, CLEC7A, CTSH, CD1C, |
| Monocyte-DC precursor | LYZ, CD1C |
| Monocyte | RNASE6, MNDA, CLEC7A, CTSH, FCN1, CD68, MS4A7, CD14 |
| Mono-NK | NKG7, CD14, GZMA, KLRB1, GZMK, STK17A, IFITM1, KLRC1, PRF1, CD3E, CD3D, CTSW, ALOX5AP, IFNG, CST7, GZMM, CD247, SH2D2A, HLA-DRB5, HLA-DPA1, HLA-DPB1, HLA-DRA, HLA-DRB1, HLA-DMA, HLA-DQB1, HLA-DQA1, HLA-DMB, HLA-DQA1, HLA-DQA2, HLA-DRB5 |
| Endothelial cell | ESAM, CD34, FGF23, FCN3, CRHBP, DNASE1L3, ACP5, RAMP2, ANGPTL4, CAV1, PRCP, TM4SF1, ECM1, KDR |
| Early Erythroid | PRSS57, GATA2, CYTL1, CLEC11A, CNRIP1, MYC |
| DC1 | BATF3, CLEC9A |
| pDC precursor | CCDC50, IRF7, IL3RA, SCT, MZB1, SPIB, IRF8, GPR183, IGLL1 |
| Mast cell | TPSAB1, CPA3, PF4, ITGA2B |
| Megakaryocyte | CMTM5, TIMP3, NRGN, GP9, PF4, PPBP, ITGA2B |
| B cell | MS4A1, CD37, CD52, TCL1A, CD79B, SPIB, FCRLA, LTB, SP140, VPREB3, BLNK, MZB1 |
| Pre B cell | IGLL1, RAG2, C1QTNF4, MS4A1, TCL1A, CD79B, SPIB, FCRLA, VPREB1, EBF1, VPREB3, BLNK, MZB1 |
| Pre pro B cell | SPINK2, IGLL1, MS4A1, VPREB1, CD79A, EBF1, MZB1, DNTT |
| Pro B cell | IGLL1, RAG1, RAG2, C1QTNF4, MS4A1, CD79B, LTB, VPREB1, EBF1, VPREB3, BLNK, MZB1, DNTT |
| Early lymphoid/T lymphocyte | STK17A, IFITM1, CD3D, CD3E, CD7 |
| NK | IFITM1, XCL2, STK17A, NKG7, KLRB1, GZMA, GZMK, KLRC1, PRF1, ALOX5AP, CLIC3, CTSW, IFNG, CD3D, CD3E, TNFSF14, CD7, CST7, GZMM, CD247, SH2D2A |
| ILC precursor | RORC, KLRB1, IFITM1, KLRB1, GZMK, CD3E, KLRC1, CTSW, CLIC3, CD247 |
| HSC/MPP | SPINK2, CD34, PRSS57, CYTL1, CLEC11A, C1QTNF4 LYZ, MPO |
| Neutrophil-myeloid progenitor | SPINK2, MPO, LYZ, CD34 |
| MEMP | CTNNBL1, PRSS57, GATA1, GATA2, CYTL1, CLEC11A |
| Erythroid | GYPA, KLF1, FAM178B , KCNH2, REXO2, CNRIP1, MYC |
| Hepatocyte | FABP1, APOA2, ALB, APOA1, SERPINA1, AHSG |
| Fibroblast | RBP1, ECM1 |
