## Supplementary Figure 1-3; Supplementary Table 1-5 for "Decoding the development of the blood and immune systems during human fetal liver haematopoiesis"

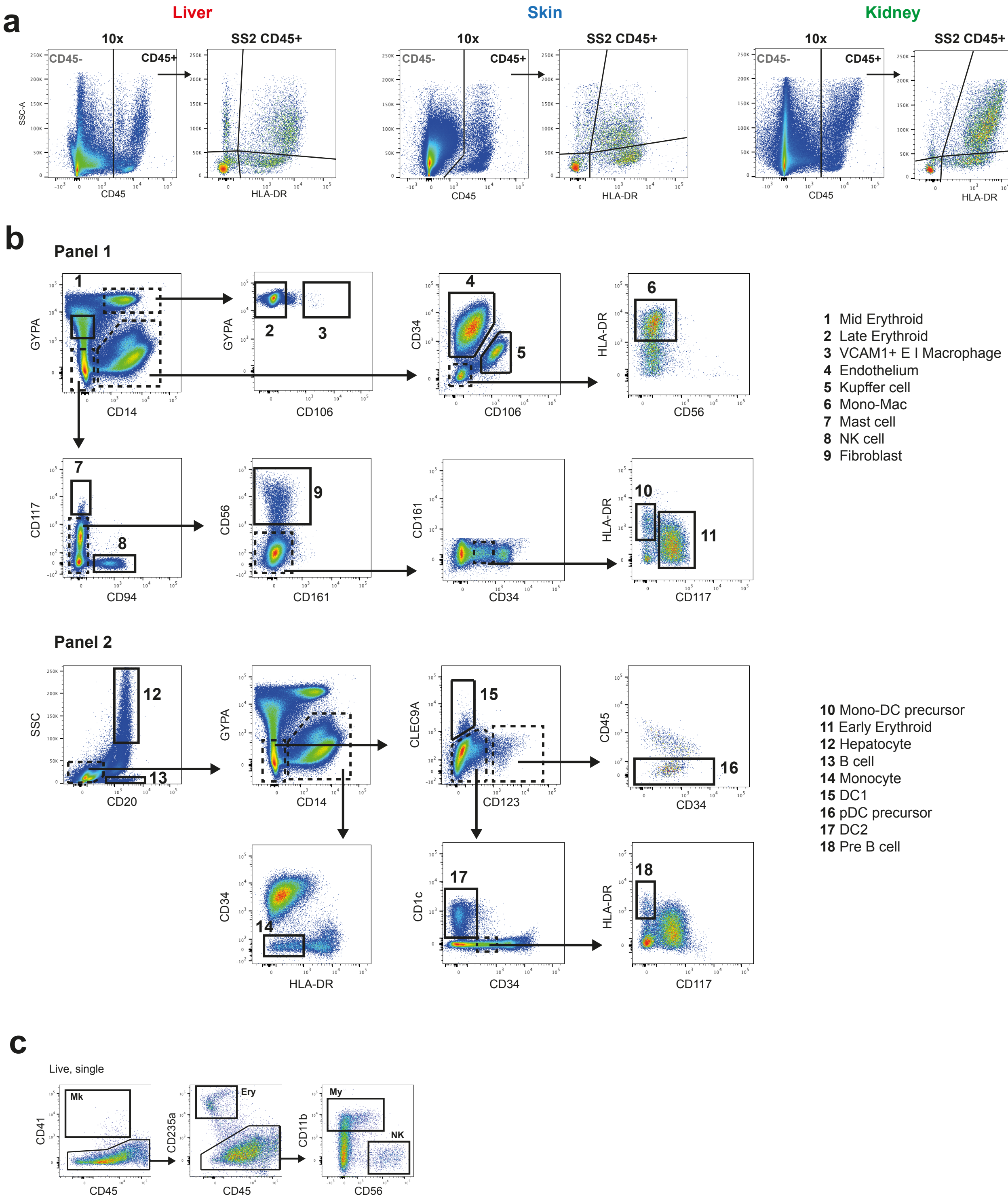

### Supplementary Figure 2

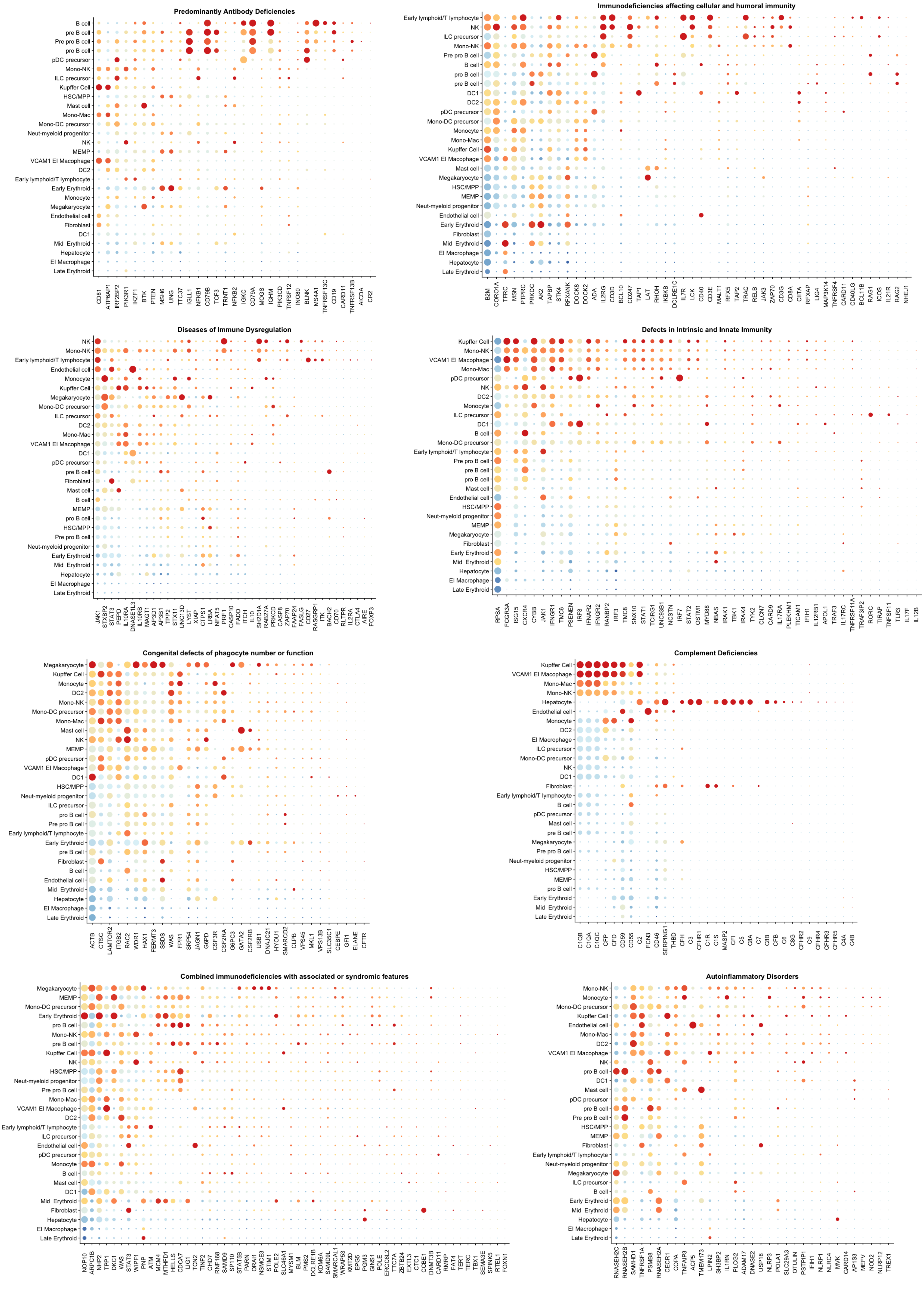

#### Supplementary Figure 3

**a**

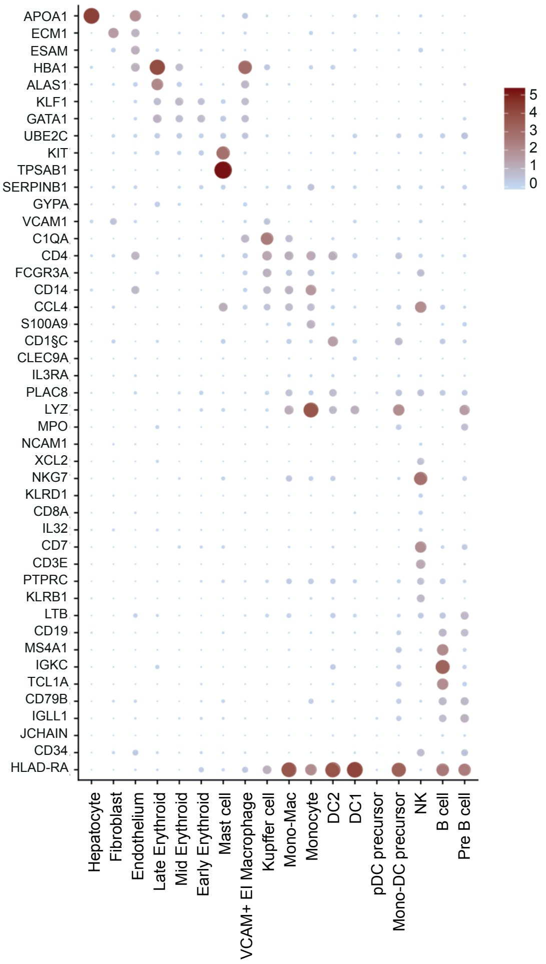


**b**

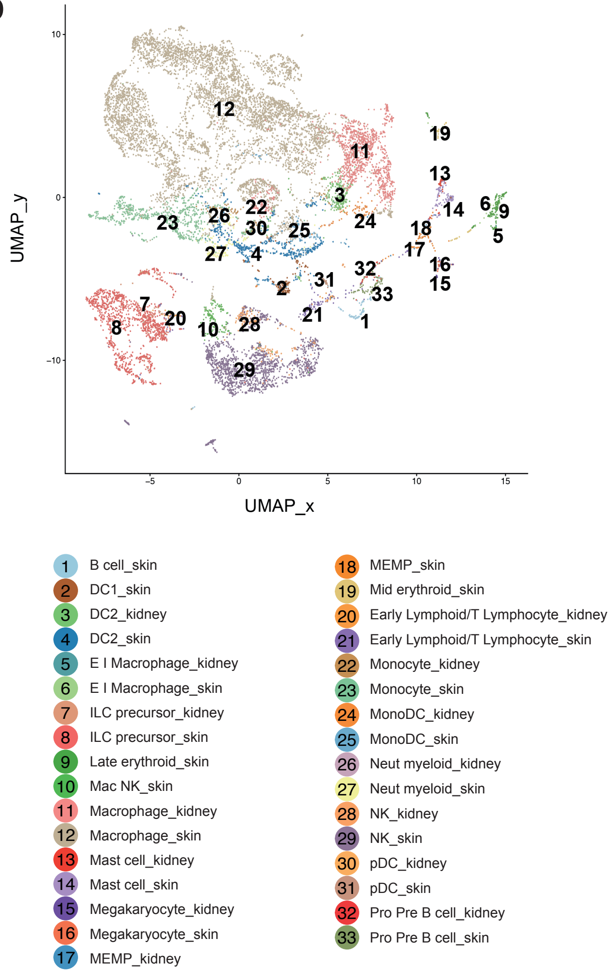

**Supplementary Table 1: Index Sort Antibodies**

| Antibody | Clone | Manufacturer |
|---|---|---|
| CD45 APC-H7 | 2D1 | BD Bioscience |
| HLA-DR BV785 | L243 | Biolegend |

**Supplementary Table 2: Cytospin/Mini Bulk Sort Antibodies**

| | Antibody | Clone | Manufacturer |
|---|---|---|---|
| **Panel 1** | CD7 BV421 | M-T701 | BD Bioscience |
| | CD14 PE CF594 | MφP9 | BD Bioscience |
| | CD34 APC-Cy7 | 581 | Biolegend |
| | CD41 AF700 | HIP8 | Biolegend |
| | CD45 BUV395 | HI30 | BD Bioscience |
| | CD56 PE | NCAM16.2 | BD Bioscience |
| | CD94 APC | REA113 | Miltenyi |
| | CD106 FITC | 51-10C9 | BD Bioscience |
| | CD117 PE-Cy7 | 104D2 | Biolegend |
| | CD161 PerCP-Cy5.5 | HP-3G10 | Biolegend |
| | CD203c BV510 | NP4D6 | Biolegend |
| | CD235a BV605 | GA-R2 | BD Bioscience |
| | HLA-DR BV785 | L243 | Biolegend |
| **Panel 2** | CD1c AF700 | L161 | Biolegend |
| | CD14 PE CF594 | MφP9 | BD Bioscience |
| | CD16 V500 | 3G8 | BD Bioscience |
| | CD19 BV421 | SJ25C1 | Biolegend |
| | CD20 FITC | L27 | BD Bioscience |
| | CD34 APC | 581 | BD Bioscience |
| | CD45 APC-H7 | 2D1 | BD Bioscience |
| | CD79A PerCP-Cy5.5 | HM47 | Biolegend |
| | CD79B PerCP-Cy5.5 | CD3-1 | Biolegend |
| | CD117 PE-Cy7 | 104D2 | Biolegend |
| | CD123 BUV395 | 7G3 | BD Bioscience |
| | CD235a BV605 | GA-R2 | BD Bioscience |
| | CLEC9A PE | 8F9 | Biolegend |
| | HLA-DR BV785 | L243 | Biolegend |

**Supplementary Table 3: HSC Culture Antibodies**

| | Antibody | Clone | Manufacturer |
|---|---|---|---|
| **Sort** | CD3 FITC | SK7 | BD Bioscience |
| | CD11c BV421 | B-Ly6 | BD Bioscience |
| | CD14 PE Dazzle | HCD14 | Biolegend |
| | CD16 FITC | NKP15 | BD Bioscience |
| | CD19 FITC | 4G7 | BD Bioscience |
| | CD34 APC-Cy7 | 581 | Biolegend |
| | CD38 PerCP-Cy5.5 | HB-7 | Biolegend |
| | CD45 BUV395 | HI30 | BD Bioscience |
| | CD45RA BV510 | HI100 | BD Bioscience |
| | CD49f PE-Cy7 | GoH3 | eBioscience |
| | CD56 FITC | NCAM16.2 | BD Bioscience |
| | CD90 APC | 5E10 | Biolegend |
| **Culture readout** | CD11b APC | ICRF44 | Biolegend |
| | CD14 APC-Cy7 | HCD14 | Biolegend |
| | CD15 BUV395 | HI98 | BD Bioscience |
| | CD41 FITC | HIP8 | Biolegend |
| | CD45 V500 | HI30 | BD Bioscience |
| | CD56 PE | NCAM16.2 | BD Bioscience |
| | CD235a BV605 | GA-R2 | BD Bioscience |

**Supplementary Table 4: IHC Antibodies**

| Antigen | Clone | Host | Manufacturer | Concentration |
|---|---|---|---|---|
| CD68 | KP1 | Mouse | Biolegend | 1:750 |
| Glycophorin A | R10 | Mouse | R&D Systems | 1:100 |

**Supplementary Table 5: Hyperion Antibodies**

| Antigen | Clone | Metal Tag | Manufacturer | Amount (ug/section) |
|---|---|---|---|---|
| CD1c | 2A7C11 | 170 Er | Novus Bio | 0.4 |
| CD20 | L26 | 141 Pr | eBioscience | 0.17 |
| CD34 | Qbend/10 | 172 Yb | Biorad | 0.8 |
| CD68 | KP1 | 153 Eu | Biolegend | 0.27 |
| CD79A | JCB117 + HM47/A9 | 161 Dy | Abcam | 0.33 |
| CK19 | polyclonal | 159 Tb | Abcam | 1.0 |
| CSP1/Heppar | OCH1E5 | 167 Er | Abcam | 0.8 |
| Glycophorin A | R10 | 142 Nd | R&D Systems | 2.0 |
